# An Open-Source Joystick Platform for Investigating Forelimb Motor Control, Auditory-Motor Integration, and Value-Based Decision-Making in Head-Fixed Mice

**DOI:** 10.1101/2025.01.23.634598

**Authors:** Ivan Linares-García, Evan A. Iliakis, Sofia E. Juliani, Alexandra N. Ramirez, Joel Woolley, Edgar Díaz-Hernández, Marc V. Fuccillo, David J. Margolis

## Abstract

Investigation of neural processes underlying motor control requires behavioral readouts that capture the richness of actions, including both categorical (choice-based) information and motor execution (kinematics). We present an open-source platform for behavioral training of head-fixed mice that combines a stationary or retractable forelimb-based joystick, sound-presentation system, capacitive lick sensor, and water reward dispenser. The setup allows for the creation of multiple behavioral paradigms, two of which are highlighted here: a two-alternative forced-choice auditory-motor discrimination paradigm, and a two-armed bandit value-based decision-making task. In the auditory-motor paradigm, mice learn to report high or low frequency tones by pushing or pulling the joystick. In the value-based paradigm, mice learn to push or pull the joystick based on the history of rewarded trials. In addition to reporting categorical choices, this setup provides a rich dataset of motor parameters that reflect components of the underlying learning and decision processes in both of these tasks. These kinematic parameters (including joystick speed and displacement, Fréchet similarity of trajectories, tortuosity, angular standard deviation, and movement vigor) provide key additional insights into the motor execution of choices that are not as readily assessed in other paradigms. The system’s flexibility of task design, joystick readout, and ease of construction represent an advance compared to currently available manipulandum tasks in mice. We provide detailed schematics for constructing the setup and protocols for behavioral training using both paradigms, with the hope that this open-source resource is readily adopted by neuroscientists interested in mechanisms of sensorimotor integration, motor control, and choice behavior.

**Significance Statement:** Behavioral paradigms for experiments in head-restrained mice are important for investigating the relationship between neural activity and behavior. However, behavioral setups are often constrained by high cost, design complexity, and implementation challenges. Here, we present an open-source platform for behavioral training of head-fixed mice using a joystick manipulandum. The setup allows for the creation of multiple behavioral paradigms, including an auditory-motor discrimination paradigm, and a value-based decision-making task. We include detailed instructions for construction and implementation of the entire open-source behavioral platform.

## Introduction

A major goal of neuroscience is to understand the relationship between neural activity and behavior. Development of sophisticated behavioral paradigms for experiments in head-restrained mice has received considerable effort because of the ability to measure and manipulate neural activity in a genetically tractable mammalian species. However, the creation of such paradigms is often constrained by high costs, design complexity, and implementation challenges. The rise of open-source approaches in neuroscience has begun to address these barriers (Burgess et al., 2017; Bollu et al., 2019; Belsey et al., 2020; Wagner et al., 2020; Manita et al., 2022; Forghani, 2023; Gordon-Fernell et al., 2023; Ozgur et al., 2023), making diverse behavioral paradigms more widely available for studying the neural basis of behavior.

Head-fixed behaviors in mice, while limited in their naturalistic scope, offer significant advantages for studying behavior in a controlled and repeatable environment (Guo et al., 2014). These setups allow researchers to precisely combine the delivery of sensory cues with the measurement of motor outputs, providing a robust framework for implementing multiple behavioral paradigms (Bjerre and Palmer, 2020). Such paradigms include Go/No-Go tasks (Guo et al., 2014; Micallef et al., 2017; Helmchen et al., 2018), two-alternative forced-choice (2AFC) tasks (Belsey et al., 2020; Burgess et al., 2017; Morandell and Huber, 2017; Estebanez et al., 2017; Gilad et al., 2018; Guo et al., 2014; Mayrhofer et al., 2013; Ozgur et al., 2023; Pan-Vazquez et al., 2024), working memory assessments (Gilad et al., 2018; Inagaki et al., 2019), and locomotion or exploration tasks (Kislin et al., 2014; Nashaat et al., 2016; Mosberger et al., 2024). The tasks utilize a range of motor outputs, including licks (Gilad et al., 2018; Guo et al., 2014; Helmchen et al., 2018; Inagaki et al., 2019; Micallef et al., 2017; Ozgur et al., 2023), reaching platforms (Estebanez et al., 2017), and floating environments (Kislin et al., 2014; Nashaat et al., 2016). In addition, manipulanda such as turning wheels (Burgess et al., 2017; Pan-Vazquez et al., 2024) and joysticks (Belsey et al., 2020; Morandell & Huber, 2017; Mosberger et al., 2024; Yang & Masmanidis, 2020) provide access to fine-grained kinematic information in a head-fixed context, allowing for detailed dissection of neural activity and effects of optogenetic manipulations. This level of control makes head-fixed paradigms with manipulanda invaluable for dissecting the relationship between neural activity and behavior.

Recent advances have demonstrated the utility of joystick manipulanda with high spatiotemporal precision in studying motor behavior (Belsey et al., 2020; Wagner et al., 2020), including reaching tasks (Bollu et al., 2019; Contreras-Lopez et al., 2023; DeWolf et al., 2024; Estebanez et al., 2017; Miri et al., 2017; Park et al., 2022), long-term motor learning (Hwang et al., 2019; Hwang et al., 2021), reinforcement learning (Panigrahi et al., 2015; Roth et al., 2024; Yttri & Dudman, 2016), motor exploration and refinement (Mosberger et al., 2024), sensory discrimination (Franco & Gourd, 2024; Hwang et al., 2017; Yang & Masmanidis 2020), and vibrotactile sensory-motor integration (Morandell and Huber, 2017; Estebanez et al., 2017). Despite these advantages, joysticks have not been widely adopted, due in part to design complexity, high costs (with notable exceptions, such as Belsey et al., 2020; Ozgur et al., 2023), and a lack of modularity. Addressing these barriers is essential for improving accessibility and promoting the widespread use of joystick-based paradigms in neuroscience.

In this work, we present an open-source joystick platform designed to provide modularity and flexibility for diverse behavioral tasks, which we demonstrate through two novel paradigms. The first is a 2AFC auditory-motor discrimination task in which mice push or pull the joystick to report different tones. The second is a value-based decision-making task that examines decision-making strategies and value-related motor output through joystick manipulation. In contrast with existing joystick-based rigs in the field (Belsey et al., 2020; Wagner et al., 2020; Ozgur et al., 2023; Mosberger et al., 2024), our setup features a fixed-base horizontal joystick with two axes of movement in the forward-backward and upward-downward directions. The setup is also compatible with a bar to restrict joystick motion to the forward-backward dimension, thus facilitating training. In addition, our joystick can be mounted on an affordable servo motor to enable joystick presentation and retraction, limiting the mouse’s interaction with the joystick to specified time windows of behavioral trials. Our joystick platform thus adds to the literature a cost-effective and versatile solution for investigating motor control and decision-making.

## Material and Methods

### Behavior rig hardware

The hardware setup can be configured for both the auditory-motor discrimination task and the value-based decision-making task, but it consists of the same basic components that can be adjusted as needed. These include Mouse Head Plates and Holder ([Janelia HHMI Head Plate and Holder] (https://hhmi.flintbox.com/technologies/c04b8f01-f188-472a-b660-368a5f8549ad)), a restraining tube (Wagner et al., 2020), a fixed or retractable joystick, a speaker, a water spout, and a licking sensor (see Fig 1A and 1F). 3D models for both the fixed and retractable joystick are available in the supplementary materials, along with a step-by-step assembly guide.

**Figure 1.**
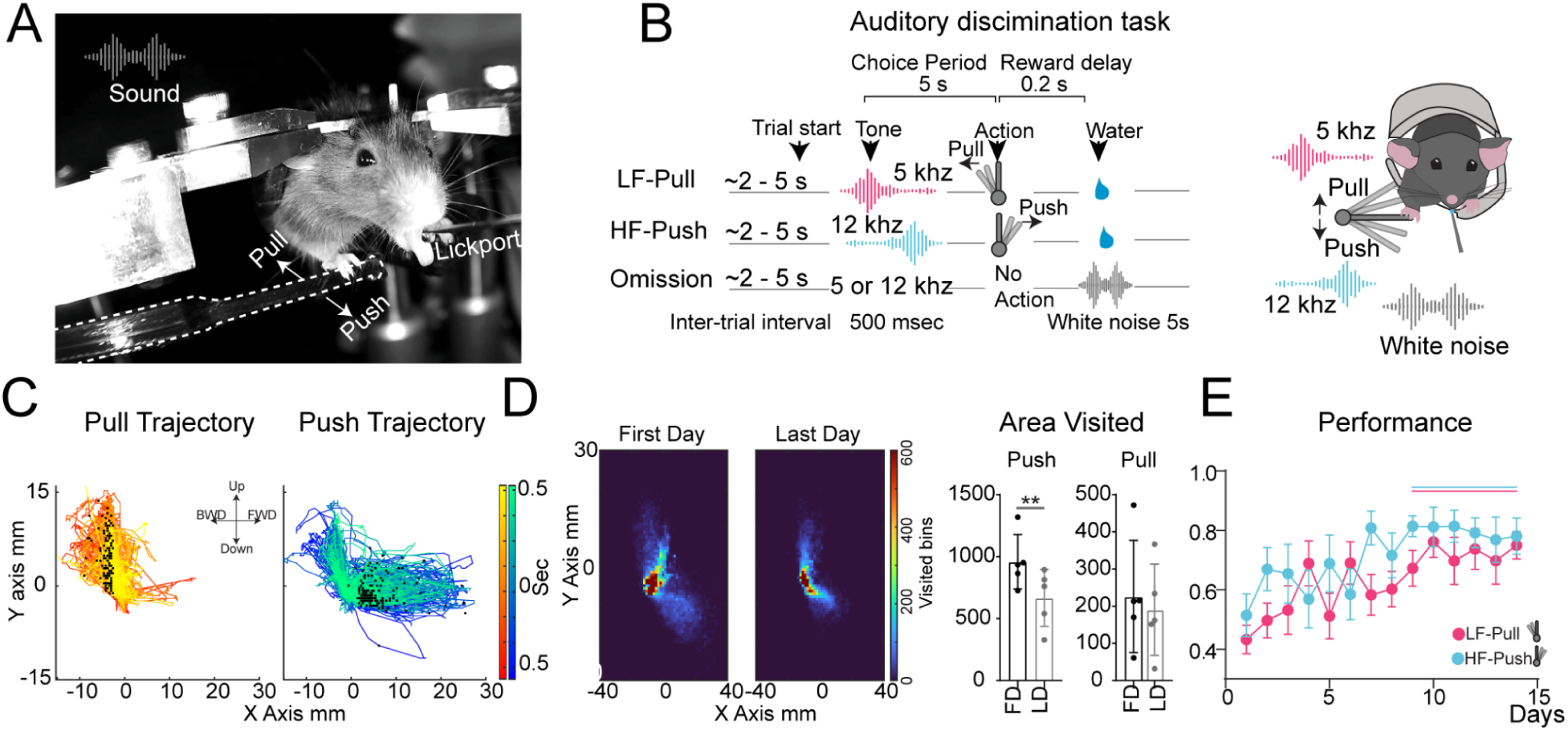
Mice learn to discriminate between distinct sounds and corresponding actions while reducing exploratory push trajectory. (A) Representative photo of an animal pushing the joystick after a tone. (B) Task schematic: mice undergo a 2–5 s inter-trial interval, then hear a 500 ms high-or low-frequency tone. Correct joystick pulls or pushes yield a water reward; omissions trigger a 5 ms white noise. (C) Joystick displacement around the choice threshold shows pulls as upward-backward and pushes as downward-forward movements. (D) Joystick area visited on first day and last day: warmer colors show higher frequency. Push area decreases with training. (E) Performance improves over time, measured as the proportion of correct trials. *n* = 10. Vertical solid lines indicate *p* < 0.05. \*\**p* < 0.0075.

The system was designed for ease of assembly, with a focus on reproducibility and scalability, allowing for creation of multiple setups within the same lab. Cooler boxes or cabinets with soundproof foam are used as chambers, while the setup is assembled using Thorlabs components, such as breadboards, optical posts, and clamps (Supplementary Material - Figures 1 and 5). This design enables modularity and flexibility to create multiple rigs—either enclosed or open—as needed for experiments involving optogenetics, calcium imaging, or fiber photometry.

The restraining tube connects to a Thorlabs clamp and an optical post, allowing height adjustments to align precisely with the head plate holders. The 3D-printed joystick system consists of two or three laser-printed parts, depending on whether the joystick is fixed in one position or retractable via a motor. The primary part is an 8 cm stick, which reduces the force required for displacement, and is glued to a 3D-printed base that secures both the stick and a 5VDC 2-axis analog APEM thumbstick for measuring motion displacements (Supplementary material Figure 1 and 6).

In the fixed joystick configuration, an additional 3D-printed part supports the joystick on one end and an adjustable friction magic arm on the other. This magic arm allows effortless positioning of the joystick, ensuring it consistently aligns below the mouse’s paw on the right side of the restraining tube. The opposite end of the magic arm attaches to an optical post mounted on a Thorlabs breadboard (Supplementary material Figure 2).

The retractable joystick setup requires two additional 3D-printed parts. The first, a reel holder, supports the joystick and enables it to slide along the second part, the servo holder, which is connected to a servo motor. This setup provides a cost-effective retractable system that can be easily controlled with a microprocessor and records joystick motion displacements (Supplementary material Figure 6).

The water spout consists of a 20G needle with a cut and smoothed tip, connected to tubing on one side of a solenoid valve (Parker: 003-0218-900 or Lee Company: LHDA1231115H) designed for silent water dispensing. The valve is connected to a tube leading to a 50 ml syringe that serves as the water reservoir. In the auditory-motor discrimination setup, the water spout is held in place by a magnetic arm and clamp attached to a steel base, which also holds a night-vision camera and a speaker (Supplementary Material - Figure 3). In the value-based decision-making setup, the water spout is integrated with an infrared lickometer (Sanworks: 1020) that is held in place by a steel arm and clamp attached to a steel base (Supplementary Material - Figure 7).

To control the auditory-motor discrimination task, we use two Arduino Uno microprocessors. The first Arduino connects to the joystick for continuous recording of motion displacements while maintaining communication with the second Arduino, which manages the entire task. Arduino 2 controls the FX sound board to play custom audio, the lick sensor through an MPR121 capacitive sensor, and the water solenoid valve via an H-bridge, allowing control of an external 12V power source. Additionally, a button is included to start the task as desired (Supplementary Material - Figure 4).

To control the value-based decision-making task, we use two Arduino microprocessors. An Arduino Mega hosts the main behavioral code, which records real-time joystick position and licks, and manages a box light, a GO cue light, and a solenoid valve (via a 12V power supply and H-bridge). It communicates with an Arduino Uno to generate pseudo-white noise via a speaker, which is used as a punishment signal (Supplementary Material - Figure 8).

This setup, while designed for the two presented tasks, can be adjusted for other task configurations by adding different components. For example, a second water port could be added for two-choice decision-making, or the sensory modality could be modified to include olfactory, visual, or whisker stimulation with minimal adjustments. This setup has also been used for calcium imaging, optogenetics, and fiber photometry (data not shown), allowing the addition of multiple transistor-transistor logic (TTL) signals to control a microscope or other devices needed for various behavioral experiments.

### Behavioral task software

#### Auditory-motor discrimination task

For the auditory-motor discrimination task, we use a primary Arduino to record joystick displacement as analog input voltage, detected through changes in resistance across two potentiometers on the X and Y axes. The Arduino’s 10-bit analog-to-digital converter (ADC) interprets analog voltage values from 0 (0 volts) to 1023 (5 volts). The joystick centers around 500 units for both axes, with displacements ranging from 9.8 mV to 24.5 mV, corresponding to changes of 2 to 5 units, and an “X” mm displacement is registered as a push or pull depending on the direction. When the threshold is reached, the primary Arduino sends a TTL signal to the secondary Arduino, which is recorded as a response from the mouse. The joystick response is sent via serial communication to be displayed and saved using a custom Python code.

The task is hardcoded on the second Arduino, which manages the sound board, joystick push/pull inputs, inter-trial interval, sound card, and the lick sensor (using a capacitive sensor). Task events are displayed and saved as a text file through the Python code. Once the programs are uploaded, both Arduinos await a TTL signal from a physical button wired to the second Arduino to start the task. This TTL signal triggers simultaneous acquisition on both Arduinos and any additional TTLs used for two-photon imaging, optogenetics, or fiber photometry.

#### Value-based decision-making task

For the value-based decision-making task, our joystick displacement readings are integrated with our Arduino Mega UART behavior code output. In addition to the joystick displacement processing steps outlined above for the auditory-motor discrimination task, we calibrate these 10-bit ADC output (0-1023) voltage readings to known displacements from baseline in millimeters on a box-by-box basis. These linear calibration functions are hardcoded into the behavior code and used to convert voltage values to millimeters in real time. We leverage the Arduino Mega’s memory and baud rate capacity to generate a 20 millisecond moving average joystick position in millimeters, to minimize contributions of aberrant spurious reads due to electrical interference. Anteroposterior deviations of greater than 3mm from baseline are registered as choices. When the joystick is retracted between trials, we generate new baseline reads to account for potential baseline drift.

This Arduino Mega also sends behavioral data to a behavior computer in real time via UART and TTL signals to our photometry and optogenetics systems. In addition, it manages our lick sensor, solenoid valve, house light, and servo motor. A separate Arduino Uno, triggered by the Arduino Mega, is used to generate pseudo-white noise as a signal that the animal has entered a time-out period following an omission or premature choice.

<u>Extended data 1.</u> (Clear instructions on how to build it)

## Results

### Mice learn to discriminate between distinct sounds and corresponding actions while refining the kinematic parameters of joystick movements

Here, we introduce a two-alternative forced-choice auditory-motor discrimination task in which animals push or pull a joystick to indicate whether they have heard a high-or low-frequency tone. This setup allows for the analysis of exploratory trajectories, velocity, tortuosity, displacement patterns, and angular motion over the course of learning, providing rich insights into motor behavior and the cognitive processes underlying decision-making.

Head-fixed water-restricted mice earn ∼10 μL water rewards by displacing a joystick in response to specific auditory cues. Joystick movements are categorized as anterior (push) or posterior (pull), corresponding to distinct high-frequency (12 kHz) or low-frequency (5 kHz) tones, respectively, each accompanied by five overtones. Reward delivery is controlled via a soundless solenoid valve equipped with a capacitive sensor at the lick spout. Auditory stimuli are presented through a front-mounted speaker controlled by a programmable sound card (Fig. 1A). The static joystick is positioned beneath the mouse’s right paw, while the left paw rests on a body tube. This setup forces right-paw use, enabling neuronal contributions to be studied through recordings or manipulations on the contralateral or ipsilateral side relative to the movement.

The task begins with a variable pre-trial interval of 2–5 seconds, followed by a 500 ms auditory cue. Mice are given a 5-second window to perform the correct joystick displacement. Correct responses trigger a 200 ms delay before reward delivery, while omissions result in a 5-second white noise signal and a reset inter-trial interval (Fig. 1B). Mice are trained in daily sessions consisting of two single-association phases: low frequency–pull and high frequency–push. Both associations are trained each day, with the training order alternating daily. Sessions last for a maximum of 30 minutes or until 100 rewards are obtained, with expert animals completing 200 correct trials and consuming up to 2 mL of water per day.

Joystick movements are recorded in two dimensions (X and Y axes), enabling the visualization of motor behavior trajectories. Push and pull actions are color-coded (red for pull, blue for push), and a temporal gradient highlights joystick movements 0.5 seconds before and after reaching the reward threshold. A black dot marks the point at which a choice was registered as either a push or a pull (Fig. 1C). Push actions involve downward-forward joystick displacement, while pull actions are characterized by upward-backward movement. This configuration provides a detailed two-dimensional representation of motor trajectories (Fig. 1C).

Exploratory behavior during learning was assessed by defining a workspace for all mice, based on the minimum and maximum x-and y-coordinates of joystick displacement across all mice. The joystick displacement workspace was binarized into smaller divisions, with each bin measuring 1 square millimeter. The trajectory areas explored were then calculated. Mice (n = 6) showed a significant reduction in the area visited during push movements between the first and last day of training (paired t-test, p < 0.0075). In contrast, no significant change was observed in the area visited during pull movements (paired t-test, p = 0.15) (Fig. 1D).

Performance was evaluated as the ratio of correct trials to the total number of correct trials and omissions. Mice exhibited significant performance improvement after session nine compared to the first day of training (two-way ANOVA, Dunnett’s multiple comparison against first day, p < 0.05) (Fig. 1E). Mice learn the auditory-motor association in 15 days, excluding 5 days of experimenter habituation during which the animals get used to handling, 2 days of head-fix habituation during which the mice freely drink water rewards while head-fixed, and 3 to 5 days of joystick association during which any displacement results in a reward, resulting in a month of training.

To quantify mouse behavior within the joystick workspace, we first identified trial-specific movement trajectories from the session data. Repeated coordinate pairs of the joystick’s location were removed, and the coordinates were centered by subtracting the median x-and y-coordinates. Each trajectory was labeled as either a push or pull trial. These trajectories were then used to calculate the average tortuosity, where higher values indicate a longer, more circuitous route from the starting position to the point of maximum displacement and back. Tortuosity was computed as the ratio of the total path length to the Euclidean distance between the first and last points in the trajectory (Mosberger et al., 2024). Session averages of tortuosity revealed that mice initially exhibited high tortuosity, which decreased and stabilized as they became proficient in the task (two-way ANOVA, Dunnett’s multiple comparison against first day, p < 0.05) (Fig. 2A).

**Figure 2.**
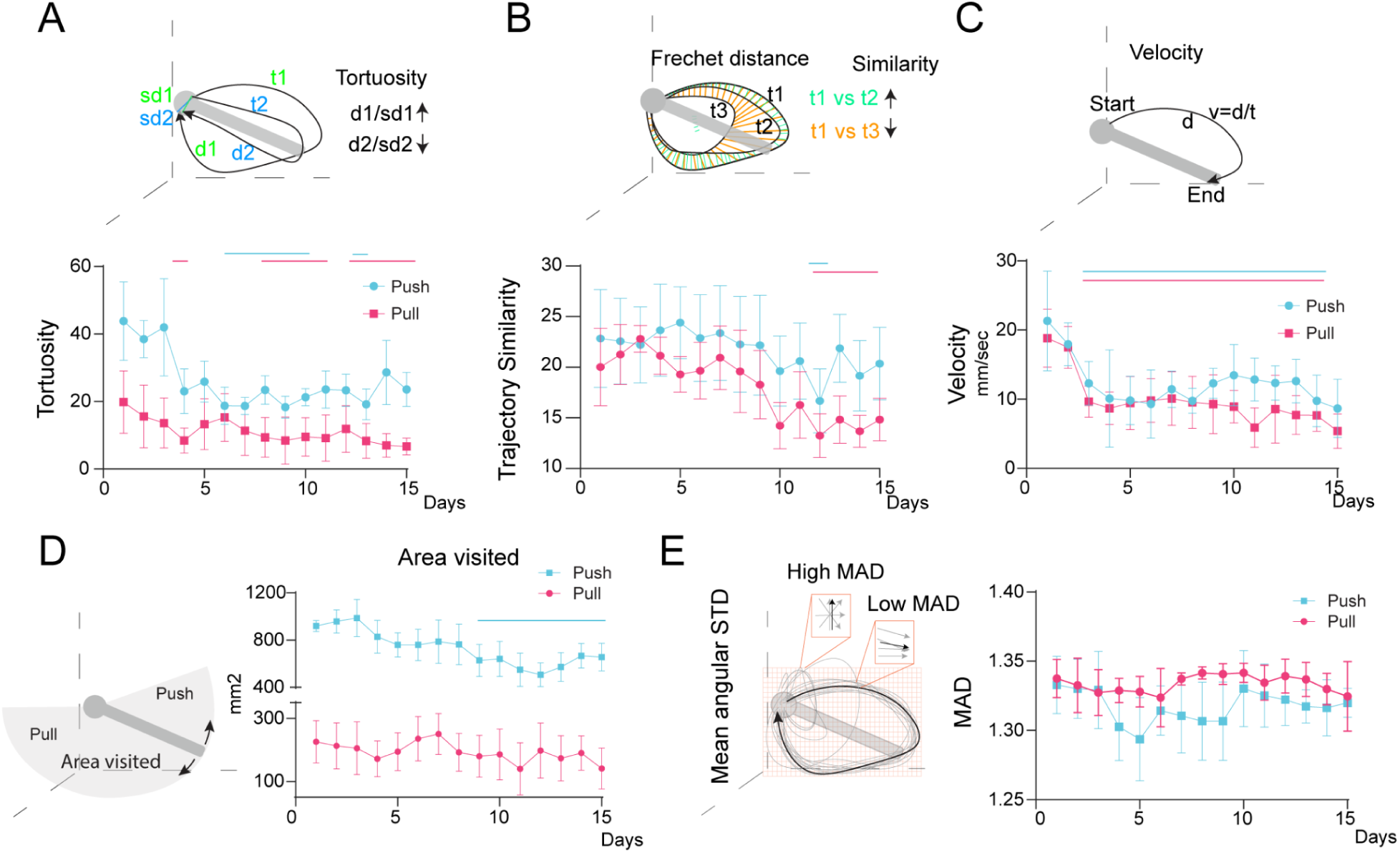
Joystick trajectory dynamics change during auditory-motor integration learning, indicating motor refinement. (A) Push and pull tortuosity across training: mice show fewer circuitous routes for both displacements. (B) Mean trajectory similarity increases across sessions, indicated by lower Fréchet distance over time. (C) Velocity to reach the displacement threshold decreases during the first days of training. (D) Joystick exploratory area visits decrease for push displacements as learning progresses. (E) Mean Angular Deviation (MAD) remains stable throughout learning. *n* = 5. Vertical solid lines indicate *p* < 0.05.

To further analyze joystick displacement dynamics, we compared the pairwise similarity of joystick trajectories using the Fréchet distance. For each session, we evaluated joystick displacement and measured the similarity across all possible combinations of trajectories, and calculated a mean value per session. Results showed that mice increased movement consistency over time, reflected by a significant reduction in the Fréchet similarity index (two-way ANOVA, Dunnett’s multiple comparison against first day, p < 0.05) (Fig. 2B).

We also measured the velocity of joystick motions, defined as the distance between the first point and the point of maximum displacement divided by the corresponding time interval. Mice demonstrated a significant increase in movement velocity compared to their performance on the first day of training (two-way ANOVA, Dunnett’s multiple comparison against first day, p < 0.05) (Fig. 2C).

To define the joystick workspace, we calculated the minimum and maximum x-and y-coordinates across all animals. The explored area within this workspace was quantified by binning the joystick coordinates using MATLAB’s “histcounts2” function, as described by Mosberger et al. (2024). Each bin measured 1 millimeter by 1 millimeter. The total explored area was calculated by summing the number of visited bins. Over successive training sessions, mice showed a significant reduction in the area explored, which eventually stabilized at a lower value, indicating reduced exploratory behavior only for the push displacement (two-way ANOVA, Dunnett’s multiple comparison against first day, p < 0.05) (Fig. 2D).

To assess directional consistency, we calculated the mean angular deviation for both push and pull motions using the CircStat MATLAB Toolbox for circular statistics (Berens, 2009). Mean angular deviation was calculated by taking the average of the angular deviation for the angles in each bin for each session. Angular deviation, ranging from 0 to √2, represents variability in directional movements, with higher values indicating greater variability. The angular deviation remained stable throughout training (Fig. 2E).

Together, these results demonstrate that mice effectively moved the joystick in two distinct directions, decreased trajectory tortuosity, increased movement similarity and velocity, and refined their displacement strategy by reducing the explored workspace. This evidence supports the task as a robust tool for analyzing motor output dynamics, offering high-quality, detailed behavioral data.

This auditory-motor discrimination task provides a robust framework for studying neural and behavioral mechanisms underlying auditory-motor associations, offering key insights into sound-driven action selection and motor learning.

#### Joystick Kinematic Parameters Reflect Total and Relative Value in Value-Based Decision-Making Task

Here, we describe a two-armed bandit, joystick-based value-based decision-making task in mice that allows for the study of value-based modulation of motor execution, in addition to recapitulating known characteristics of conventional value-based 2-AFCs.

Head-fixed water-restricted mice obtain 10% sucrose solution rewards via anterior or posterior displacement of a retractable joystick. Reward is delivered via an optical lickometer setup that also tracks licking (Fig. 3A). At trial start, the joystick is made available to the mouse via anterior motion of the servo motor. Following a subsequent 100ms wait period, a GO cue light on the lickometer signals the start of a 10-second window during which the mouse can register a choice via anterior (push) or posterior (pull) displacement of the joystick (Fig. 3B). There are four possible outcomes of a trial: (1) *rewarded trial*, followed by 2.5-8-second inter-trial interval; (2) *unrewarded trial*, signaled by turning off house light, followed by 2.5-8-second inter-trial interval; (3) *omission*, signaled by white noise, turning off house light, and 15-second time-out; (4) *premature trial* in which mouse registers choice before Go cue, signaled by white noise, turning off house light, and 15-second time-out.

**Figure 3.**
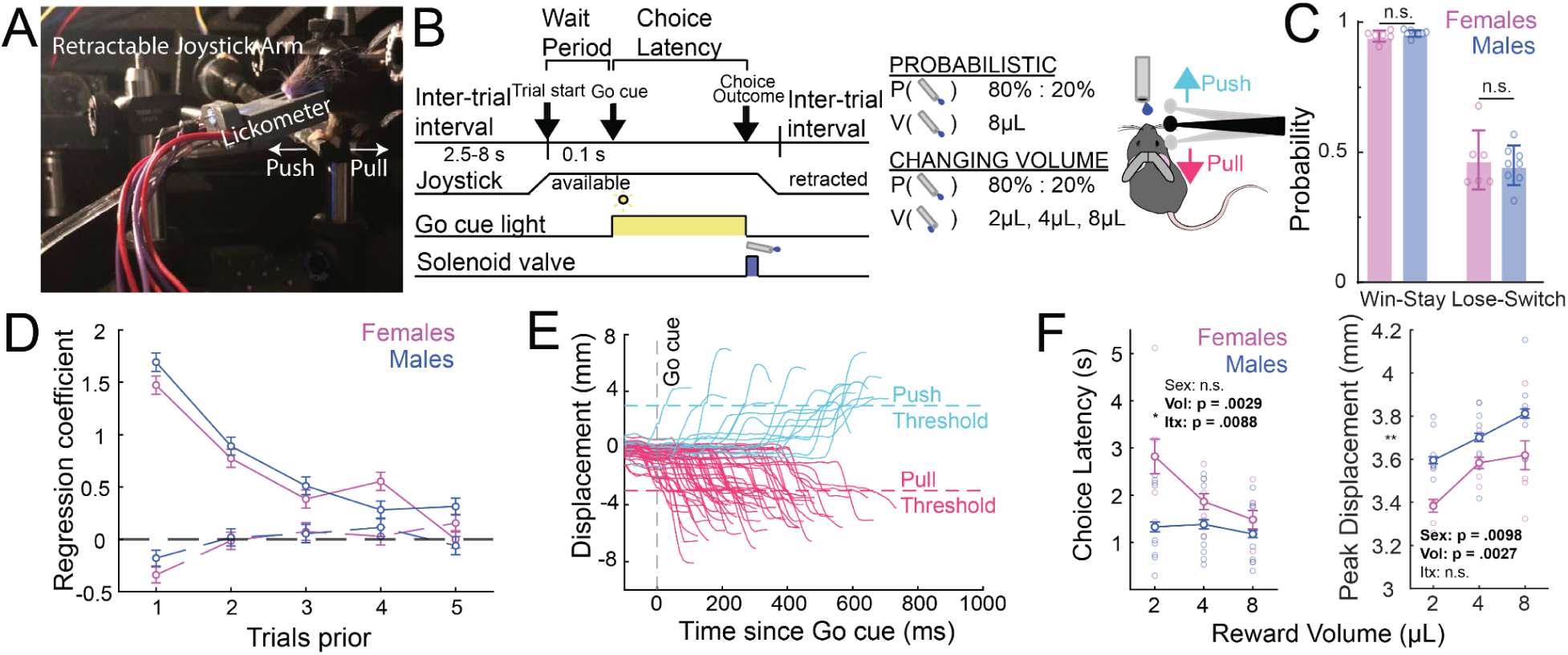
Mice integrate reward evidence across trials to guide next-trial choice and motor vigor. (A) Photograph of our behavioral setup. (B) behavioral schematic highlighting two task variants. (C) Males (n = 8) and females (n = 6) prior outcome to guide next trial strategy. (D) Logistic regression weights of prior trial rewarded (undashed) and unrewarded (dashed) outcomes for predicting current trial repetition of prior choice. (E) Raw joystick choice data. (F) Demonstration that varying reward volume significantly shapes motor vigor, as measured by peak joystick displacement and, predominantly in females, choice latency.

Mice are trained in a sequential behavioral paradigm consisting of, (1) probabilistic reversal, and (2) changing volume phases (Fig. 2B). The *probabilistic reversal* phase consists of blocks in which one of two choices is more likely to be rewarded than the other (*push* blocks and *pull* blocks), with reward probabilities of 80%:20%. Each block has a minimum duration of 17 *rewarded* trials plus a geometrically-distributed random variable (*p* = 0.4), after which the high reward probability side is reversed in an un-cued manner. As in the auditory-motor discrimination task, mice only register choices with their right forepaw. Mice in this phase integrate prior-trial evidence to guide next-trial decisions as evidenced by win-stay/lose-switch analysis as well as logistic regression of choice and reward history (Fig. 3C,D). The *changing volume* phase builds on this task structure by adding reward volume as an additional parameter that varies by block, using reward volumes of 2, 4, and 8 μL. Mice register choices with varying latencies and peak joystick displacement (Fig. 3E). We find that, in higher total value contexts, mice register choices with shorter choice latency and higher peak joystick displacement (Fig. 3F). Unlike choice latency, the peak displacement phenotype is robust across males and females, suggesting that a joystick-based design offers unique, key insights into animals’ regulation of motor vigor based on total value as compared to conventional, binarized lever press-or lick-based tasks (e.g., Alabi et al., 2020, Wang et al., 2013).

An advantage of our joystick-based value-based decision-making paradigm over conventional lever-press or lick-based paradigms in mice is the ability to read out kinematic parameters of operant choice. Vigor is known to reflect real-time internal value representations (Shadmehr et al., 2019; Takikawa et al., 2002).

In this task, we find that mice’s joystick trajectories often exhibit extensive deliberation before crossing choice threshold (Fig. 4A). Antero-posterior joystick position relative to baseline can be plotted as a function of time and segmented into movement bouts to capture these ***deliberative*** movement bouts leading up to a threshold-crossing ***decisive*** bout. We defined bouts according to the following criteria: (1) movement is in one direction, (2) initiation speed is greater than 7.5mm/s, (3) speed is maintained at >2.5mm/s for >50ms, (4) bout ends with velocity sign change or joystick retracting. Of note: unlike the auditory-motor discrimination task, our value-based decision-making task constrains motion to the antero-posterior axis via addition of a metal bar underneath the joystick, limiting up-down joystick displacement.

A range of kinematic parameters can be extracted from joystick movement traces. ***Peak displacement*** is defined as the maximum extent of displacement of the joystick away from baseline on a given trial. ***Number of bouts*** is defined as the number of movement bouts the animal initiates in a given trial. ***Directional consistency*** is the proportion of these bouts that occur in the higher frequency direction (Equation 1), with a value of 1 implying that all bouts occur in one direction. ***Mean velocity*** is defined as the mean velocity of the joystick on the decisive bout. ***Tortuosity*** is defined mathematically as the ratio of (1) the distance travelled by the joystick in any direction on a given trial and (2) the displacement of the joystick on that trial from baseline to peak (peak displacement). When mice move the joystick straight from the center to threshold, the distance travelled by the joystick should be close to the displacement, yielding a tortuosity value of close to 1. If the path of the joystick is more winding (i.e., tortuous), the distance travelled by the joystick is much greater than the end-to-end displacement, yielding a higher tortuosity value.

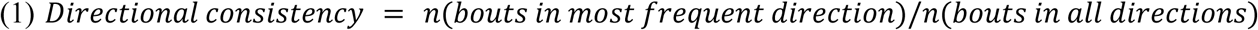

We found that our readouts of joystick trajectory reflect animals’ internal representations of certainty that one joystick direction is more likely to yield reward than another, i.e., relative value. Using a two-parameter Q-learning model with nondifferential forgetting (Ito & Doya, 2015; J Neurosci; Choi et al., 2023; Nat Commun), we generated trial-by-trial estimates of animals’ internal representation of the value of *push* and *pull* actions (Q_push_ and Q_pull_). We computed the absolute value of the difference between Q_push_ and Q_pull_ (ΔQ = Q_push_-Q_pull_) to gauge the animals’ experienced uncertainty on a given trial, where low |ΔQ| implies more similar value representations of push and pull actions, and therefore greater uncertainty regarding which choice is more likely to be rewarded. Movement trajectories in high |ΔQ| (low uncertainty) and low |ΔQ| (high uncertainty) contexts are distinct, as is illustrated in example traces (Fig 4B). In lower |ΔQ| trials, we found that animals trended towards lower peak displacement (Fig. 4C), and had significantly greater mean joystick velocity (Fig. 4D), a significantly greater number of joystick movement bouts (Fig. 4E), significantly lower directional consistency (Fig. 4F), and significantly greater tortuosity (Fig. 4G), reflective of greater uncertainty. Our joystick kinematic parameters thus provided a key additional insight into animals’ dynamic representations of relative value.

**Figure 4.**
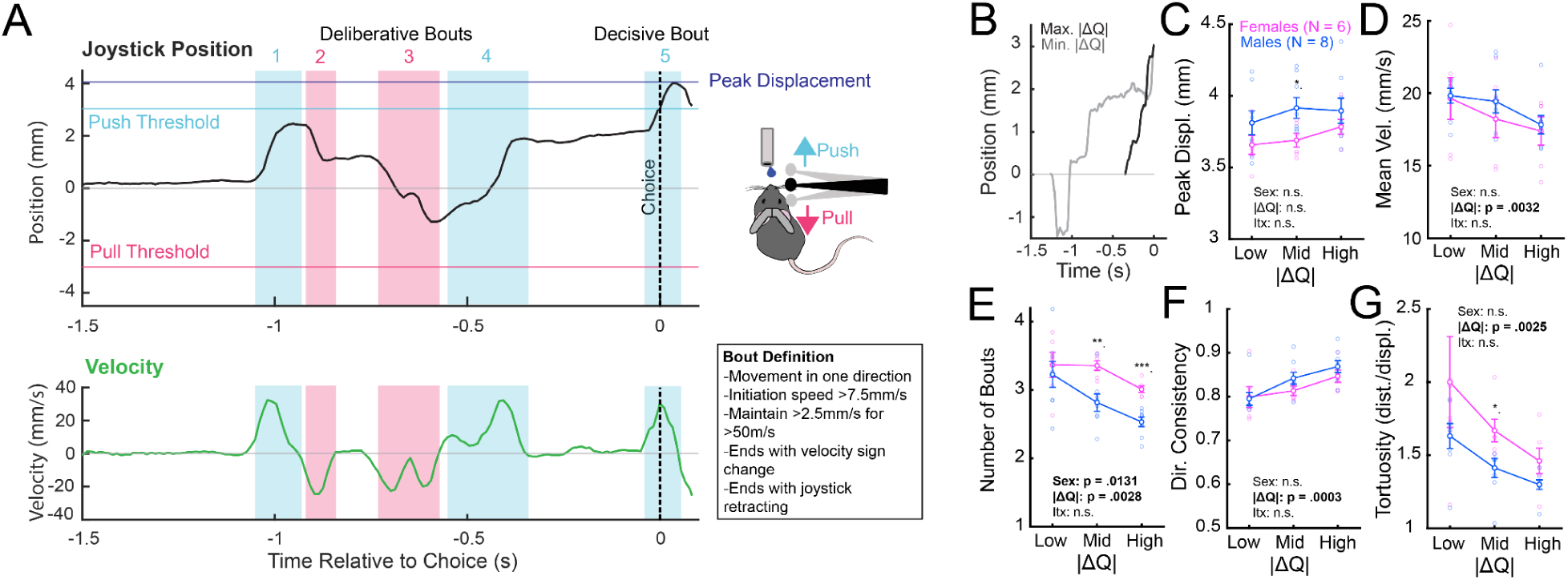
Readouts of joystick trajectory in different relative value contexts reflect animal uncertainty. (A) Segmentation of joystick position trace into movement ‘bouts’ based on joystick velocity. (B) Representative traces from high uncertainty (low |ΔQ|) and low uncertainty (high |ΔQ|) contexts. In higher-uncertainty contexts, mice register choices with greater mean velocity (D), greater number of movement bouts (E), lower consistency of direction of joystick motion across bouts (F), and higher tortuosity (G). Peak joystick displacement is not significantly affected by degree of uncertainty (C). * p <0.05; ** p < 0.01; *** p < 0.001.

## Discussion

Our work documents an open-source platform for behavioral training of head-fixed mice using a forelimb-based joystick manipulandum. We demonstrate, in a novel 2AFC auditory-motor discrimination paradigm, that mice refine their motor execution throughout learning. Similarly, as mice learned a novel value-based decision-making task, they constrained their motor vigor in the context of lower reward volume and uncertainty. In these tasks, the joystick provided kinematic readouts reflective of learning stage and internal value representations. Joystick kinematic parameters thus provided key additional insights into the motor execution of choices that are not as readily assessed in freely moving, lever press, or head-fixed licking contexts.

Disentangling sensory neuronal encoding from motor output is complex yet crucial for decision-making (Mohan et al., 2018; Ranieri et al., 2022). While some brain areas are primarily involved in sensory processing, others, including the striatum, serve as integrative hubs for both sensory and motor inputs (Gerfen, 1984; Hunnicut et al., 2016), with individual neurons often receiving both sensory-and motor-related synaptic input (Lee et al., 2019; Sanabria et al., 2024; Ramanathan et al., 2002; Assous and Tepper, 2019). In this context, paradigms that disentangle sensory inputs from motor outputs are essential for identifying their distinct contributions to neuronal activity (Morandell and Huber, 2017; Burgess et al., 2017; Estebanez et al., 2017; Ozgur et al., 2023). While simpler sensory discrimination paradigms, such as Go/NoGo licking tasks, can confirm an animal’s ability to distinguish stimuli, they provide limited insight into the decision-making process beyond sensory discrimination (Guo et al., 2014). 2AFC behavioral paradigms can be used to explore perceptual decision-making (Bjerre and Palmer, 2020). By presenting two stimuli and associating them with two distinct actions, these paradigms allow for the generation of different behavioral metrics to compare and contrast against neuronal activity. This approach facilitates testing for selectivity and distinguishing between sensory stimulus, choice, motor action, and outcome selection (Chen, Susu, et al., 2024). Here, we introduce a 2AFC auditory-motor discrimination paradigm that incorporates custom sounds—tones with overtones at high and low frequencies. Mice are trained to discriminate between these sounds and report their choices through distinct joystick movements: anterior (push) or posterior (pull) displacements executed with a single forepaw. Our paradigm could be modified easily to include other sensory cues (e.g., visual, tactile, olfactory) relevant for investigating sensory discrimination and cued movements in multiple modalities.

A joystick-based task design also offers significant advantages in the study of value-based decision-making. In addition to recapitulating aspects of known features of two-armed bandit designs, including integration of evidence across trials and adaptation of behavior as contingencies change (Chantranupong et al., 2023; Parker et al., 2016; Tai et al., 2012), it enables the study of value-related invigoration of movements as is seen classically with saccades in primate value-based decision-making designs (Reppert et al., 2015; Shadmehr et al., 2019; Takikawa et al., 2002). We demonstrate that our joystick kinematic metrics, such as peak displacement, mean velocity, tortuosity, and properties of movement bouts, are reflective of animals’ internal total and relative value representations, as captured by standard reinforcement learning algorithms. Given the intricate interplay of value-and vigor-related information in cortex, basal ganglia, and the midbrain (Dudman & Krakauer, 2016; Nakamura & Hikosaka, 2006; Hikosaka et al., 2014; Niv et al., 2006; Shadmehr et al., 2019; Wang et al., 2013), this task provides rich behavioral outputs with which to study the neural representation of value-based decision-making in mice.

We present two distinct behavioral paradigms built on a shared hardware design, offering a versatile framework adaptable to diverse experimental needs. These paradigms can be modified to accommodate different sensory modalities by altering the stimuli. For example, whisker stimulation can be implemented using a 12V stepper motor paired with an Adafruit motor shield for Arduino and 3D-printed windmill textures. Similarly, visual stimulation can be introduced using an Adafruit SSD1327 OLED Graphic Display interfaced with Arduino via I2C. Additionally, a simple sensory discrimination paradigm can be incorporated through optogenetic stimulation of sensory inputs triggered by Arduino transistor-transistor logic signals (TTLs; Sachidhanandam et al., 2013). The H-bridge used to drive the water solenoid is designed to support an additional solenoid. This feature enables the integration of a second water spout, facilitating the development of a head-fixed version of a two-step task (e.g., Thomas Akam et al., 2021) or devaluation paradigms (e.g., Turner & Balleine, 2023). Because our setup operates using Arduinos, it can easily interface with fiber photometry, optogenetics, or 1/2-photon calcium imaging via TTLs, facilitating study of the neural basis of behavior.

A limitation with head-fixed, joystick-based setups is their relative difficulty. Head-fixation per se can delay learning timelines as it can increase animal anxiety and is less ‘naturalistic’. In addition, while mice readily learn to displace the joystick manipulandum within a couple of days, it is anecdotally more difficult for mice to learn to distinguish two different directions. This part of training requires attention and can take up to a month, as is also seen in other joystick-based paradigms (e.g., Hwang et al., 2021). One way to expedite training is to constrain joystick motion using bars to minimize out-of-plane motion, or force the mouse to move the joystick in a non-preferred direction (i.e., forcing a mouse to pull that prefers to push). Another potential limitation is that our joystick comes in from the right side, and cannot be displaced along the left-right axis. It is therefore not ideal in the study of left vs. right choice as is seen in some basal ganglia studies (e.g., Tai et al., 2012, Bolkan et al., 2022), which would require left-right joystick designs (e.g., Belsey et al., 2020). Where the kinematics of action execution are not of interest, head-fixed licking-based paradigms or freely moving lever/nose-poke based paradigms should be preferred as these might be more readily learnable.

Investigation of the neural processes underlying motor control requires precise behavioral readouts that capture the kinematics of motor actions. Here, we present a low-cost, open-source, joystick-based platform for the behavioral training of head-fixed mice, which allows for the study of learning and task-related refinement in motor execution. The joystick metrics we highlight provide only a glimpse into the wealth of spatiotemporal data that can be extracted from our real-time joystick position recordings. We hope this setup will be readily adopted and expanded upon by the neuroscience community to provide insights into the kinematic parameters of sensorimotor integration, decision-making, value representation, and other neural processes.

## Acknowledgements

This work was supported by grants from the NIH (F30-MH136699 to E.A.I., R01-MH118369 to M.V.F., R01-NS094450 to D.J.M.) and NSF (IOS-1845355 to D.J.M.). J.W. was supported by UPenn NIH Training Grant T32-NS105607. I.L.-G. was supported by a Rutgers Busch Biomedical Research Grant. We thank Thomas J. Vajtay for assistance with 3D designs and hardware development, Dr. Alex Yonk and members of the Margolis lab for useful discussions. We thank Sarah Ferrigno for her advice in task design and training, and Luigim Vargas-Cifuentes for his assistance with our reinforcement learning model design. We also thank Alessandro Jean-Louis and Wenxin Tu for excellent technical assistance.

## Author Contributions

Conceptualization - I.L.-G., E.A.I., M.V.F., D.J.M.

Methodology - I.L.-G., E.A.I., E.D.-H., M.V.F., D.J.M.

Formal analysis - I.L.-G., E.A.I., S.E.J., J.W.

Investigation - I.L.-G., E.A.I., A.N.R.

Writing - Original Draft - I.L.-G., E.A.I., M.V.F., D.J.M

Writing - Review and Editing - I.L.-G., S.E.J., E.A.I., A.N.R., E.D.-H., M.V.F., D.J.M

Funding Acquisition - I.L.-G., E.A.I., J.W., M.V.F., D.J.M.

## Supplementary material - Linares-García, Iliakis, et al

### Part 1. Auditory-Motor discrimination task Head Fix setup assembly

To assemble the Head-Fix setup, you will use a 3D-printed body (as described by Wagner et al., 2020), a tube, and the Janelia HHMI head post assembly. Follow these detailed instructions to securely attach these components to a breadboard using pedestal posts and clamps.

1. **Breadboard Base Setup**:

a. Use a Thorlabs breadboard (MB2020/M) as the base.
b. Assemble three pedestal posts using TR4 optical posts and insert them into PH50/M post holders.
c. Attach BE1/M pedestal adapters at the base of each post.
d. Secure the posts to the breadboard using CF125C/M clamps, positioning the first clamp at C2 and the second at C7. Use M6 10mm screws at positions E2 and E7, and tighten them with a 3/16” hex key.
2. **Head Mount Bar and Post Holder Assembly**:

a. At the top of the optical posts, secure the head mount bar from Janelia Designs using an M4 16mm screw in the second hole, tightening it with a 7/64” hex key.
b. Insert the bottom piece of the headpost holder into the sliding hole of the head mount bar near the end closest to the adjacent bar. Secure it using an M4 16mm screw, a 7/64” hex key, and an M4 hex nut.
c. Attach the top part of the headpost holder to the bottom piece using an 8-32 x 1/4 cap screw. Repeat this for the opposite bar.
d. Ensure the correct distance between the bars by fitting a head plate into the headpost holder holes.
3. **Securing the Third Post**:

a. Position the third post using a CF038 clamp at B5, near the right-side bar.
b. Attach the RA90/M right-angle post clamp at the top of this post and secure it with a hex key.
4. **3D-Printed Body Tube Installation**:

a. Attach the 3D-printed body tube to the TR20/M post.
b. Insert the body tube into the second hole of the RA90/M clamp and secure it.
c. Adjust the height of the body tube so it sits between the two bars

**Figure 1.**
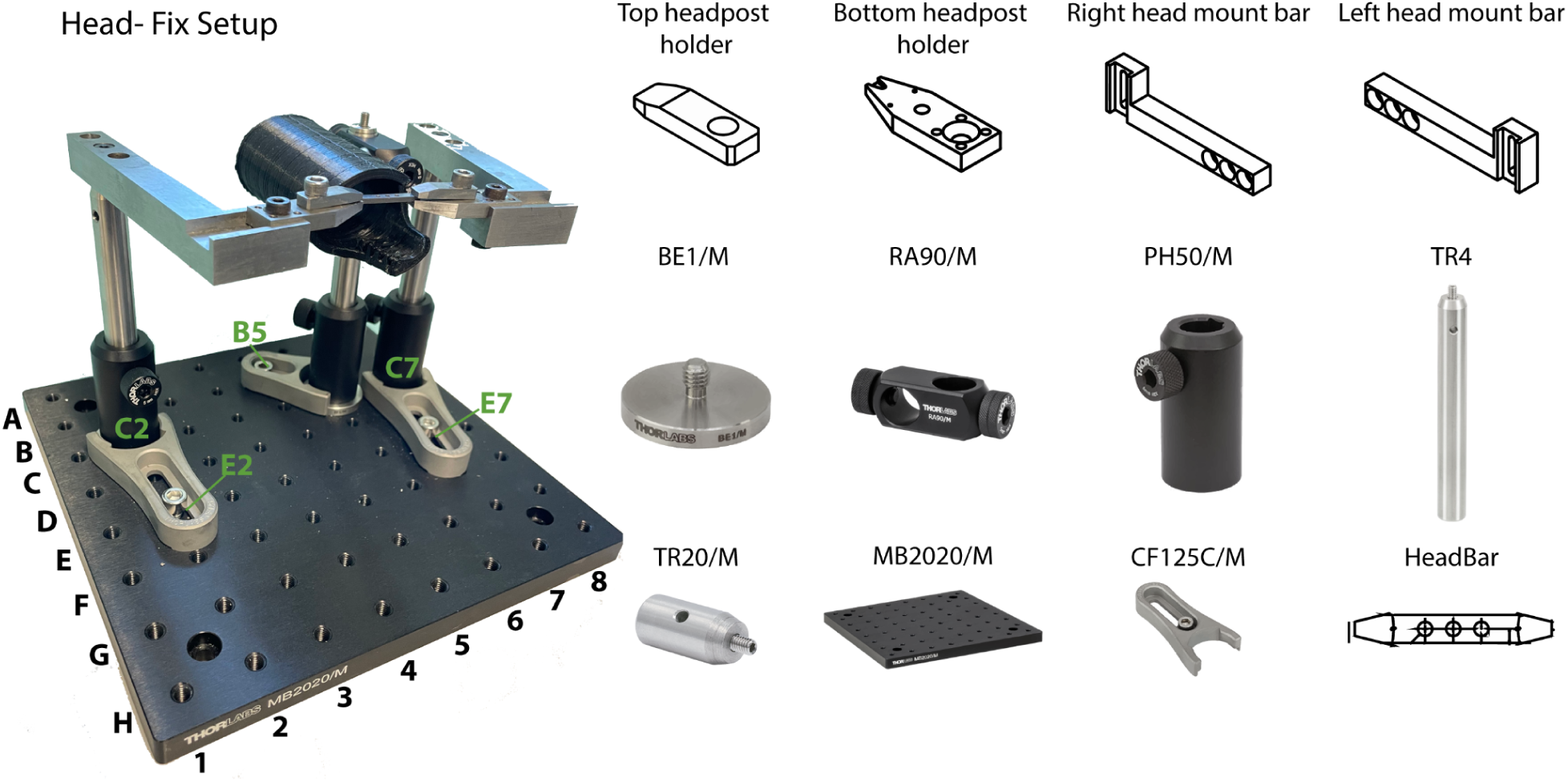
**Auditory discrimination task Head Fix setup assembly.**

### Fixed Joystick setup assembly

To assemble the fixed joystick, begin by attaching the thumbstick to a 3D-printed assembly that consists of three main components: the stick lever, stick base, and joystick base holder. These parts will be mounted on a magic arm and a breadboard, allowing you to position the joystick comfortably for the mice to achieve optimal displacement.

1. **Breadboard Base Setup**:

a. Use the Thorlabs breadboard (MB1015/M) as the base for mounting.
b. Assemble one pedestal post using the TR4 optical post and insert it upside down into the PH50/M post holder.
c. Secure the PH50/M post holder to the breadboard at the C1 position using an M6 x 25 mm cap screw.
2. **Magic Arm Assembly**:

a. Attach one end of the magic arm to the port on the PH50/M post holder and secure it firmly.
b. On the other end of the magic arm, attach a TR20/M post and secure it.
3. **Joystick Assembly**:

a. Attach the joystick base holder to the TR20/M post using an M4 16mm screw.
b. Insert the joystick into the base holder hole so that the joystick extends outside the hole and base.
c. Secure the joystick with the hex nut that comes with it on the opposite side of the hole.
d. Remove the black plastic casing surrounding the joystick.
4. **Stick Base and Lever Installation**:

a. Apply T-7000 adhesive to the inside part of the stick base, then glue it securely to the joystick.
b. Once the adhesive is set and the stick base is firmly attached, insert the stick lever into the stick base and secure it with a screw if needed.

This assembly will ensure that the joystick is stable and accurately positioned for use.

**Figure 2.**
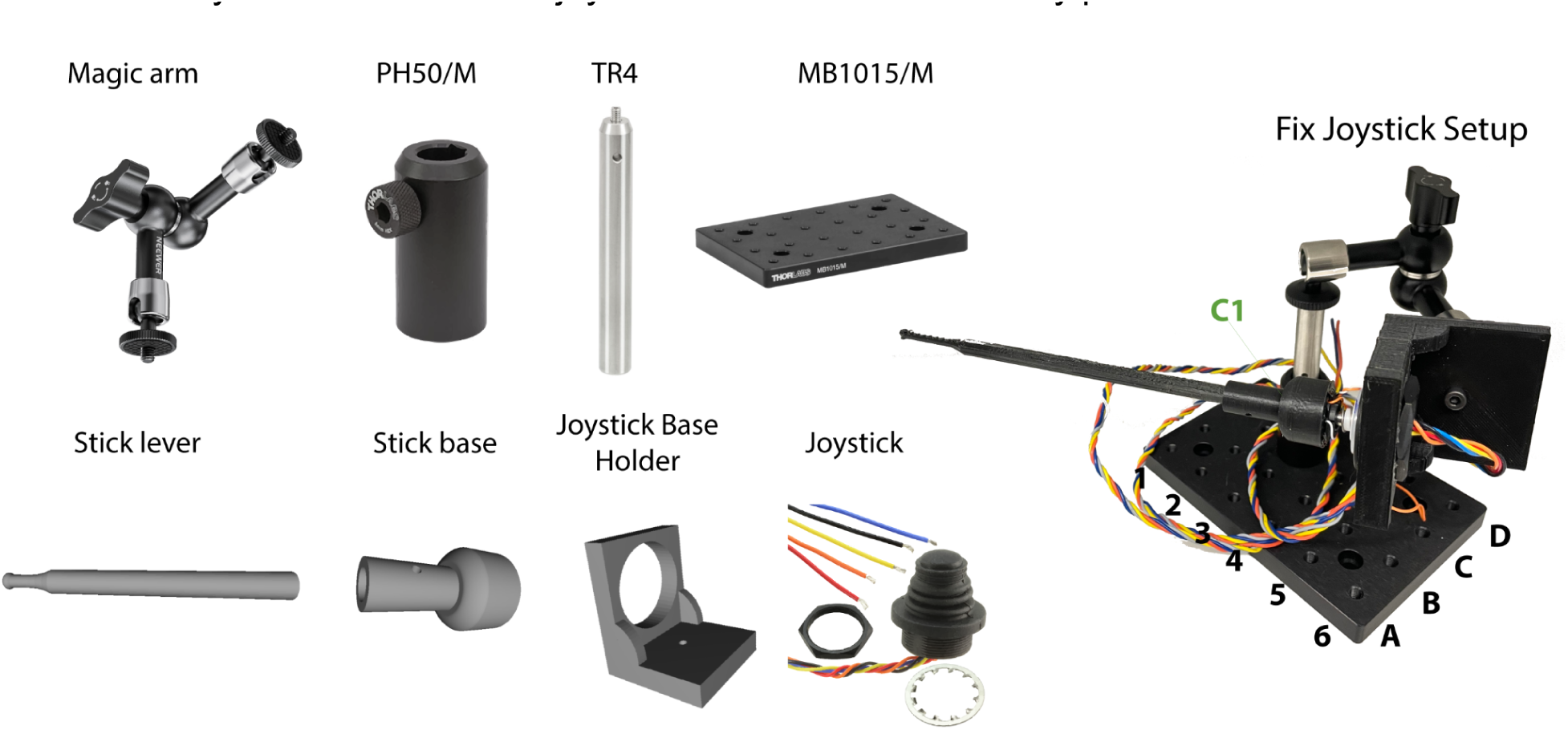
**Fixed Joystick setup assembly**

### Speaker, camera and water spout setup assembly

To assemble the complete behavioral box, you will integrate the previously built joystick assembly and head-fix setup along with a speaker, camera, and water spout. These components will be installed inside a cooler that will serve as a soundproof behavioral chamber.

1. **Cooler Preparation**:

a. Remove the plastic water leak plug from the cooler to expose a hole that will connect the inside of the box (where the behavioral setup is located) to the outside (where the microprocessors will be housed).
b. Line the sides, front, and top of the cooler with soundproof foam. Use T-7000 glue to adhere the foam to the cooler’s interior. Let the foam set and rest overnight to eliminate any lingering odors.
2. **Integrating Joystick and Head-Fix Setup**:

a. Join the joystick and head-fix setups by aligning the breadboard bases. Secure them together using M6 10mm screws at the D2 and B2 positions.
b. Adjust the magic arm to position the joystick close to the gap in the 3D-printed body tube, as shown in the reference image.
3. **Mounting the Speaker, Camera, and Water Spout**:

a. Use a steel base to mount the speaker, camera, and water spout assembly.
b. Detach the clamp from two magic arms and remove the screw at the tip. Insert the magic arms into the top two holes of the speaker and tighten the screws to secure it.
c. For the lick port, use another magic arm to hold the needle that will be attached to the solenoid. Attach a wire to the metal part of the needle, which will be connected to a capacitor sensor to detect licks.
d. Use an additional magic arm to mount the camera, ensuring it is positioned to record the mice during the experiment.
e. This Infrared camera allows to record mice in the absence of light therefore no additional illumination is needed in the box to train animals.

This setup will allow you to create a functional and soundproof environment for behavioral experiments. For the electronic diagram on how to connect the hardware follow next instructions.

**Figure 3.**
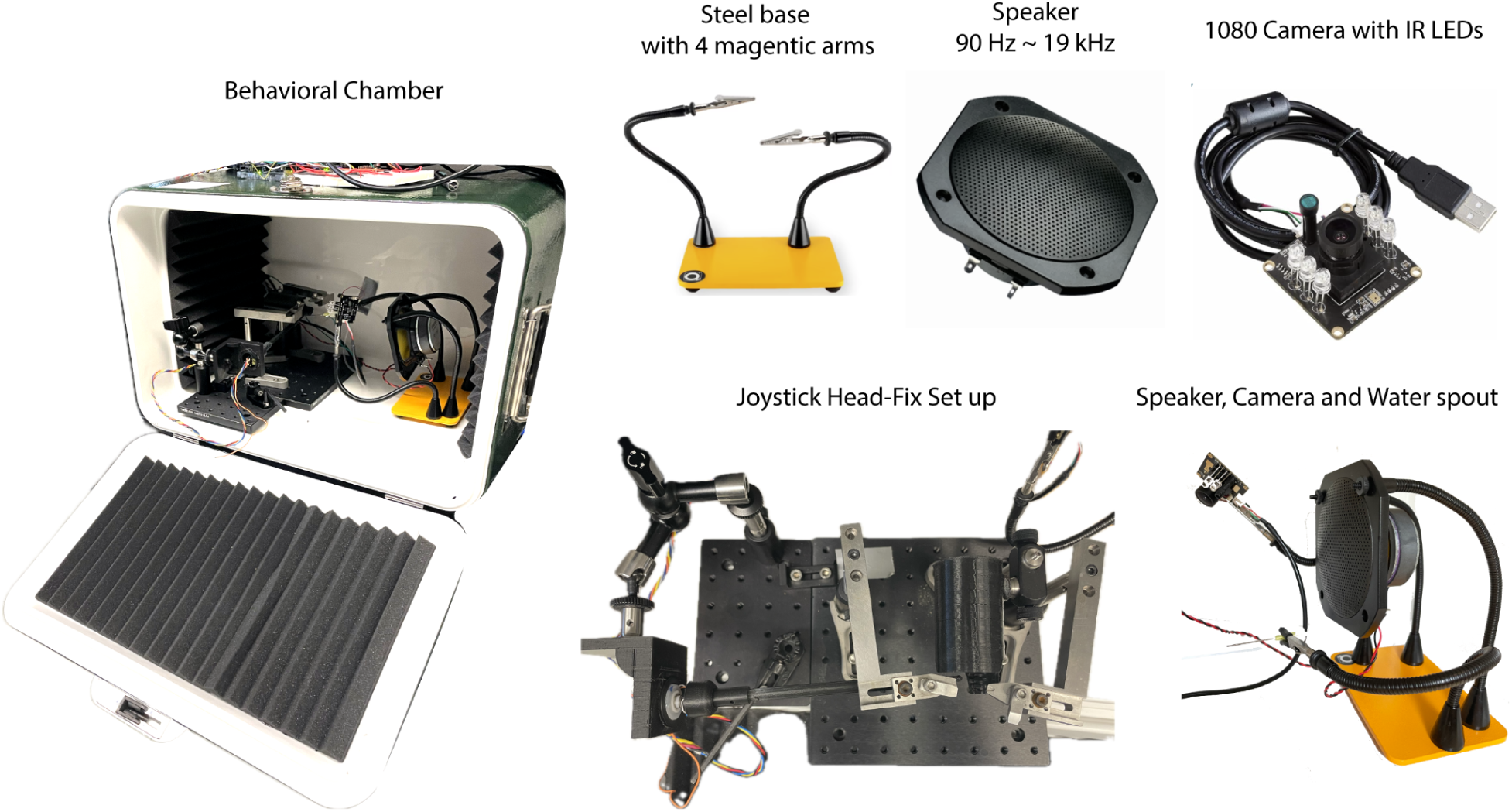
**Speaker, camera and water spout setup assembly**

### Electronic Diagram

To assemble the electronic diagram for the dynamic auditory action selection paradigm, follow these steps to connect the components, including two Arduinos, the joystick, an FX Sound Board, an MPR121 capacitive sensor, and a Parker Series 3 noiseless solenoid valve.

1. **Joystick Connection to Arduino 1**:

a. Connect the joystick wires to Arduino 1, which will handle continuous joystick displacement data collection.
b. Wire the joystick as follows:

i. Red voltage cable to 5V input on Arduino 1.
ii. Black cable to ground (GND).
iii. Blue cable to analog input 1 (A1).
iv. Yellow cable to analog input 3.
c. Since the joystick cables are short, extend them using additional wires and soldering, ensuring a length of at least two meters for mobility.
2. **Interfacing Arduino 1 and Arduino 2**:

a. Maintain continuous communication between Arduino 1 and Arduino 2 by connecting the following pins:

i. Pin 13 on Arduino 1 to A0 on Arduino 2.
ii. Pin 12 on Arduino 1 to A1 on Arduino 2.
iii. Pin 11 on Arduino 1 to pin 12 on Arduino 2.
iv. Pin 8 on Arduino 1 to pin 3 on Arduino 2.
v. Pin 7 on Arduino 1 to pin 8 on Arduino 2.
3. **Arduino 2 and Breadboard Power Distribution**:

a. Connect the 5V input from Arduino 2 to the left voltage column of the first breadboard.
b. Connect the 3.3V input to the second left voltage column.
c. Connect VIN to the right voltage column of the third breadboard.
d. Ground (GND) from Arduino 2 should be connected to the left side of the first breadboard. All breadboards will share the same ground from Arduino 2.
4. **FX Sound Board Setup**:

a. Prepare the sound card for breadboard mounting by soldering the provided pins. Position it from row one to row fourteen.
b. Connect the following wires:

i. Ground (GND) from Arduino 2 to sound card ground.
ii. UG pin on the sound card to Arduino 2 GND.
iii. RX to pin 6 on Arduino 2.
iv. TX to pin 5 on Arduino 2.
v. Vin to the 5V input on Arduino 2.
vi. Voltage and ground wires from the speaker (previously soldered).
5. **MPR121 Capacitive Sensor Integration**:

a. Solder the pins on the MPR121 touch sensor and mount it on a second breadboard next to the first one, spanning rows one to thirteen.
b. Connect the following:

i. 3.3V from Arduino 2 to 3Vo on the touch sensor.
ii. GND to GND.
iii. A5 to SCL.
iv. A4 to SOA.
v. Pin 11 on Arduino 2 to INT.
6. **Push Button Installation**:

a. Place a push button between the eighteenth and twentieth rows of the second breadboard, with the middle part of the breadboard separating the rows.
b. Connect the following:

i. Eighteenth row (left side) to input 2 on both Arduino 1 and Arduino 2.
ii. Eighteenth row (right side) to Arduino 2 GND using a 1kΩ resistor.
iii. Twentieth row (right side) to Arduino 2 5V.
7. **H-Bridge and Solenoid Wiring**:

a. Attach a third breadboard next to the second one. Place the H-bridge in the middle, spanning rows one to eight.
b. Connect both sides of row one to the 5V input on Arduino 2.
c. Connect pin 7 of Arduino 2 to the second row of the H-bridge.
d. Connect the fourth row (left side) to Arduino 2 ground.
e. Connect the eighth row (right side) to the VIN pin on Arduino 2, which will receive 15 volts from an external adapter.
f. Connect the two wires from the solenoid to the third and sixth rows of the H-bridge, with the order of connection not being crucial.

This assembly ensures that the dynamic auditory action selection paradigm is fully operational with all components integrated and connected for precise task management and data collection.

**Figure 4.**
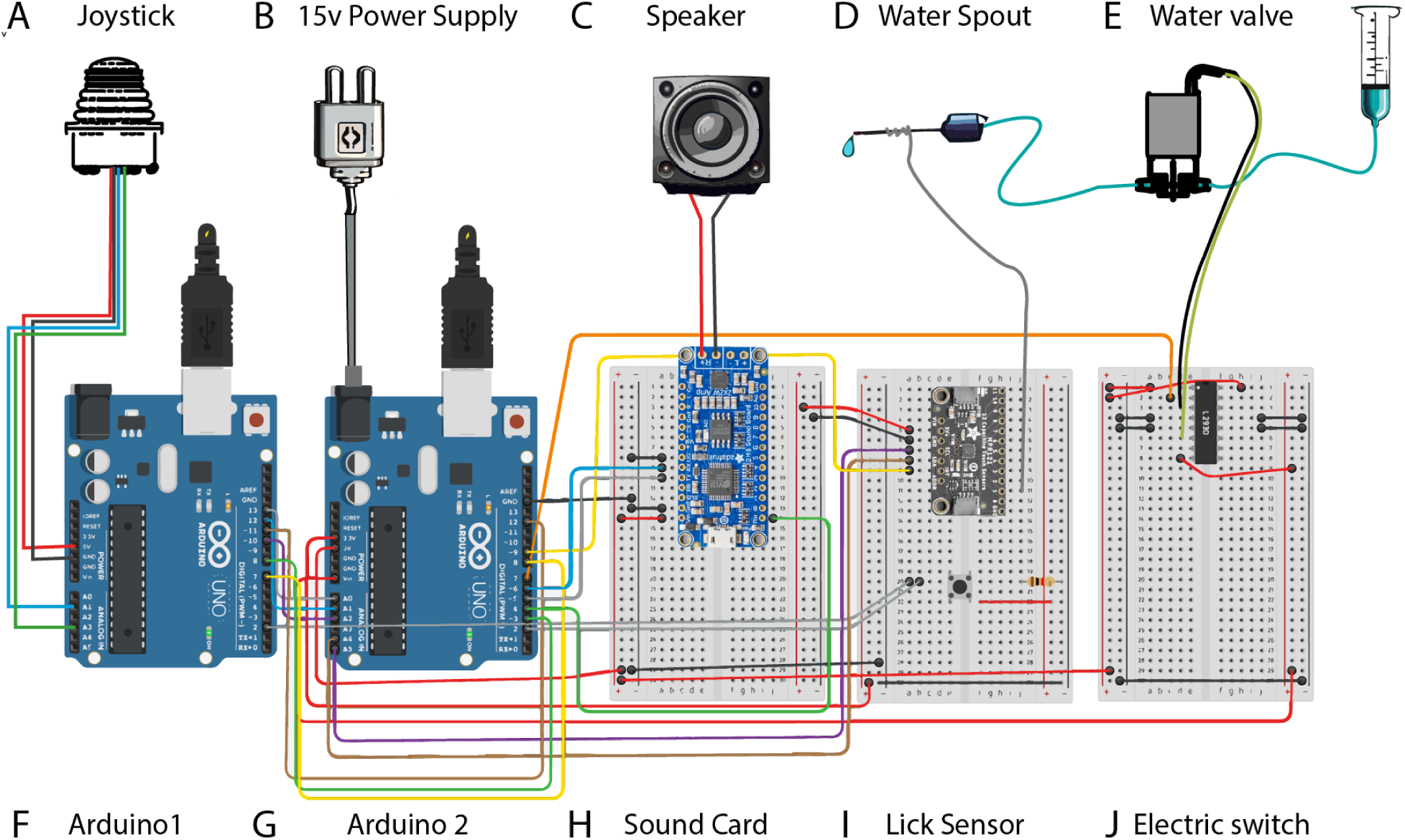
**Auditory discrimination task electronic diagram.**

### Part 2 - Value-Based Decision-Making Task Rig Assembly

Note: this setup is all built on a single breadboard (Thorlabs MB1012 Aluminium Breadboard). The holes in this breadboard are arranged in a 10-by-12 configuration. In these instructions, we will refer to these holes using the following coordinate system:

**Figure 5.**
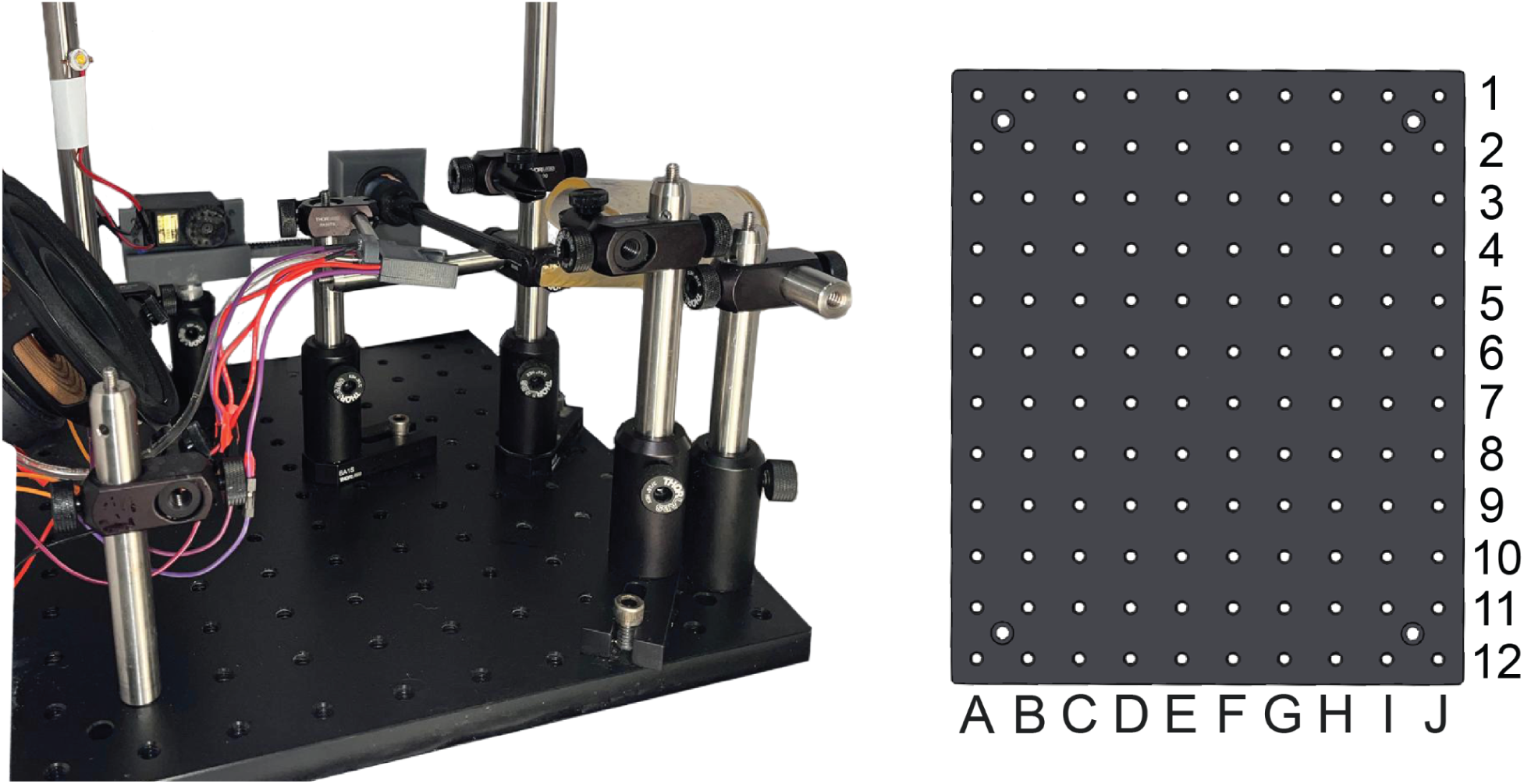
**Value-based decision-making task rig assembly.**

### Gluing steps

1. Use epoxy resin to glue a 2” post (TR2) to the laser-printed motor holder. Ensure that the motor holder is as level as possible, as this step will determine the angle at which the anteroposterior movement of the joystick occurs. Let dry overnight.
2. To generate the wheel that will extend and retract the joystick, cut a piece of timing belt (GT2) long enough to loop around the wheel included in the servo motor (SG-5010, Adafruit: 155), and attach it using epoxy resin. Let dry overnight. When attaching, ensure that the timer belt overhangs in the same direction that the wheel will interface with the servo motor. If it overhangs in the opposite direction, it will not fit in the motor holder.
3. To generate the interface of the joystick reel with the servo wheel, cut a piece of timing belt (GT2) long enough to cover the entire long and thin portion of the joystick wheel. Stick to the joystick reel with epoxy resin, and let dry overnight.
4. To generate the base of the joystick, remove the plastic cover of the thumbstick (TS6T2S02A, DigiKey: 679-3658-ND) and fix the joystick holder to the exposed tip of the thumbstick (TS6T2S02A, DigiKey: 679-3658-ND) using epoxy resin, and let dry overnight. Applying epoxy resin minimally and carefully to avoid disrupting the seamless mobility of the thumbstick: avoid the spring and the plastic plates closest to the joystick base.

### Assembly steps

1. Using 4 screws (SH3M10), screw the servo motor (SG-5010, Adafruit: 155) onto the motor holder glued to the 2” post (TR2).
2. Attach the wheel from the servo motor (SG-5010, Adafruit: 155) glued to the timer belt (GT2) to the servo motor (SG-5010, Adafruit: 155) at the white shaft interface. Ensure it snaps into place.
3. Insert the joystick onto the joystick base glued to the thumbstick (TS6T2S02A, DigiKey: 679-3658-ND). Some plastic might need to be scraped off the joystick to ensure a snug fit. In the case of a loose fit, some epoxy resin is recommended to stabilize the joystick.
4. Insert the rear end of the thumbstick (TS6T2S02A, DigiKey: 679-3658-ND) into the joystick reel, facing away from the long thin part of the reel, and screw tight with the hex nut included with the thumbstick (TS6T2S02A, DigiKey: 679-3658-ND). **It is critical at this juncture to ensure that the white and orange wires at the base of the thumbstick are perfectly aligned to the top/12-o’clock position of the joystick holder as shown in the image, as this will ensure the correct orientation and magnitude of the XY displacement readouts of the joystick.**
5. Guide the assembled joystick reel glued to the timer belt (GT2) into the servo motor holder at the space between the motor wheel and the bottom of the motor holder, until the tip reaches the level shown on the image above. Note that this level can and should be adjusted to ensure optimal positioning of the joystick relative to the mouse.
6. Screw a post holder (PH2) into the coordinate B4 of the breadboard (MB1012) using screw SS8S050.
7. Insert the 2” post glued to the motor holder into the post holder (PH2) and screw to tighten.

**Figure 6.**
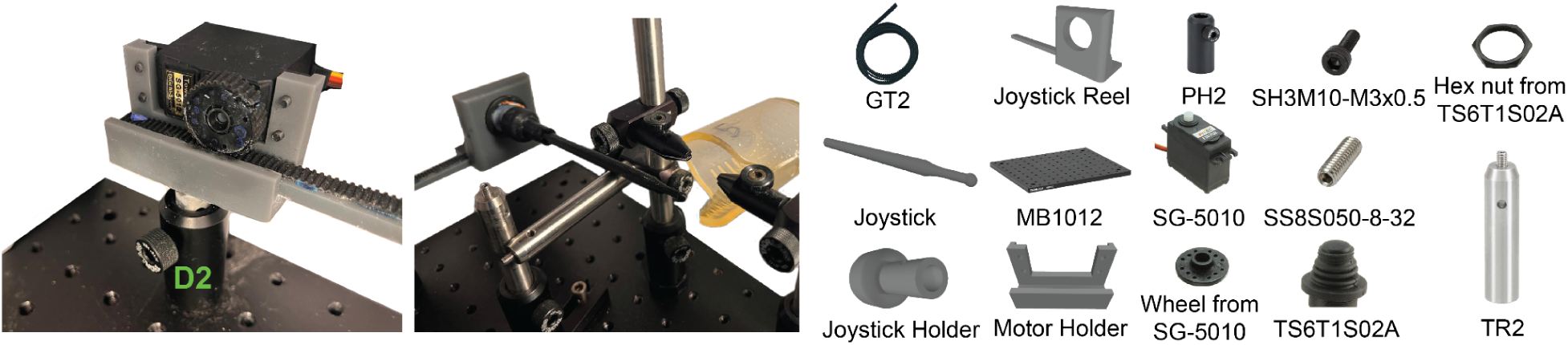
**Retractable joystick assembly.**

### Head-Fix Setup and Lickometer Assembly

#### Set up Mouse Tube

1. Screw post holder (PH2) into breadboard at coordinate I12 using screw SS8S050.
2. Insert 4” post (TR4) into the post holder at I12. Tighten using the built-in horizontal screw on PH2.
3. Slide right-angle clamp (RA90) onto the 4” post and tighten onto post using the appropriate screw on the RA90.
4. Insert an 8-32” hex nut into its corresponding opening in the laser-printed mouse tube.
5. Tightly screw the mouse tube onto a 2” post (TR2), by screwing the small built-in screw on the post onto the 8-32” hex nut through the hole in the mouse tube.
6. Insert the TR2 post with the attached mouse tube into the horizontal hole of the RA90 clamp attached to the 4” post on I12. This tube will be used to hold the mouse as it performs the behavioral task.
7. Adjust height and position of mouse tube so that the tip of the extended joystick is at the level shown below:

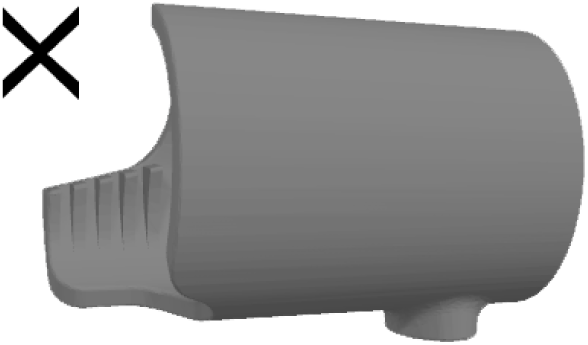

#### Assemble Proximal Head Clamp

1. Screw a post holder (PH2) to mounting base (BA1S) using screw SH25S075V.
2. Insert 4” post (TR4) into this post holder (PH2), tighten using horizontal screw on post holder (PH2).
3. Slide right-angle clamp (RA90) onto the 4” post (TR4) and tighten onto post using the appropriate screw on the RA90.
4. Insert post clamp (PC2) into the horizontal hole of the right-angle clamp (RA90). Tighten the right-angle clamp (RA90) screw to fix the post clamp (PC2) in place. This clamp will be used to hold the side of the mouse’s headplate closest to the outside of the box (proximal to the experimenter).

#### Assemble Distal Head Clamp

1. Screw a post holder (PH2) to mounting base (BA1S) using screw SH25S075V.
2. Insert 4” post (TR4) into this post holder (PH2), tighten using horizontal screw on post holder (PH2).
3. Slide right-angle clamp (RA90) onto the 4” post (TR4) and tighten onto post using the appropriate screw on the RA90.
4. Insert 4” post (TR4) into horizontal hole of this right-angle clamp (RA90). This post will be adjusted so it sits right underneath the joystick to prevent mice from displacing the joystick downwards.
5. Slide a second right-angle clamp (RA90) onto the 4” post (TR4) and tighten onto post ∼5cm above the first right-angle clamp using the appropriate screw on the RA90.
6. Insert post clamp (PC2) into the horizontal hole of this second right-angle clamp (RA90). Tighten the right-angle clamp (RA90) screw to fix the post clamp (PC2) in place. This clamp will be used to hold the side of the mouse’s headplate closest to the inside of the box (distal to the experimenter).

#### Position Head Clamps Around Mouse Tube

It is recommended to position the head clamps around the mouse tube at the height of the top of the mouse tube, using the configuration shown below:

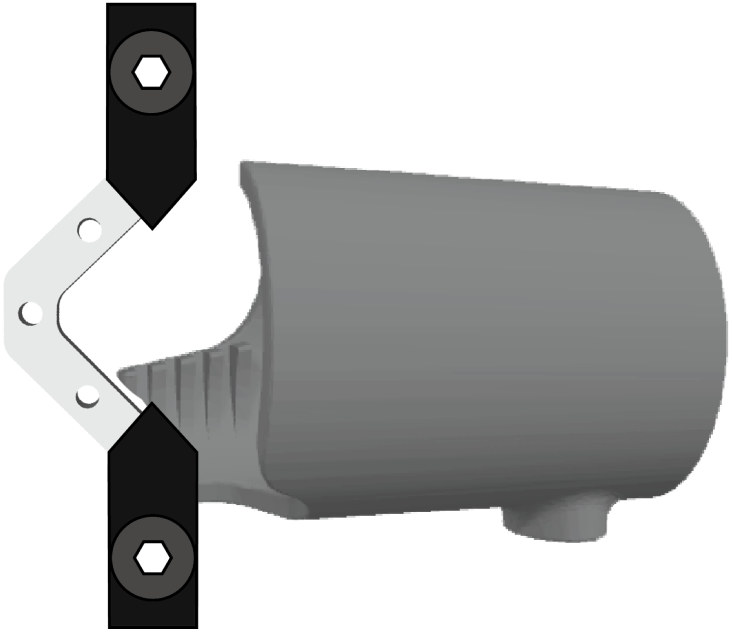

In our setup, this position is accomplished most easily by screwing in the distal clamp setup at D12 on the breadboard, and the proximal clamp setup at J10, using screws SH25S075V.

#### Assemble Lickometer Setup

1. Mount lickometer onto mini-series optical post (MS3R) by screwing small attached screw mini-series optical post (MS3R) into small hex nut built into the lickometer. Set aside.
2. Screw a post holder (PH2) to mounting base (BA1S) using screw SH25S075V.
3. Insert 4” post (TR4) into this post holder (PH2), tighten using horizontal screw on post holder (PH2).
4. Slide **1”-1/2“** right-angle clamp (RA90**<u>R</u>**) onto the 4” post (TR4) and tighten onto post using the appropriate screw on the RA90R.
5. If needed, loosen small hex screw on RA90R, then slide on MS3R attached to lickometer. Tighten the small hex screw on the RA90R to have a somewhat firm grip on the lickometer, allowing for some slack to tilt the lickometer up and down, and move it towards and away from the post, but keeping it tight enough to prevent the lickometer from tilting down on its own.
6. Position the lickometer setup in front of the mouse tube, around 2.5cm below the height of the clamps. This is best achieved by screwing the mounting base (BA1S) into hole D9 of the breadboard.

**Figure 7.**
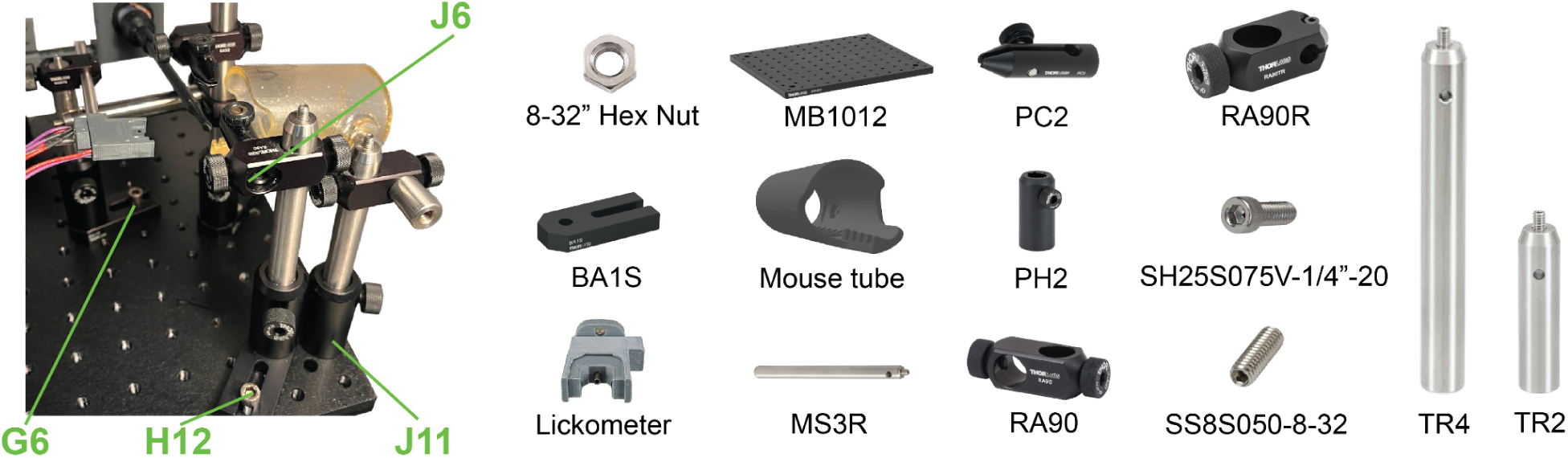
**Head-Fix Setup Assembly and Lickometer Assembly**

#### Speaker Assembly

1. Screw the 8” post (TR8) into hole C2 of the breadboard (MB1012), and the 4” post (TR4) into hole I2 of the breadboard (MB1012) using screw SS8S050.
2. Slide a right-angle clamp (RA90) onto each post as shown in the image.
3. Insert a post clamp (PC2) into each of the horizontal holes of the right-angle clamp (RA90) and tighten them in place using the hex screw of the right-angle clamp (RA90).
4. Insert the bottom two corners of the speaker into the post clamps (PC2) and tighten until stable.
5. Attach box LED to top of the 8” post (TR8) to serve as a box light.

**Figure 8.**
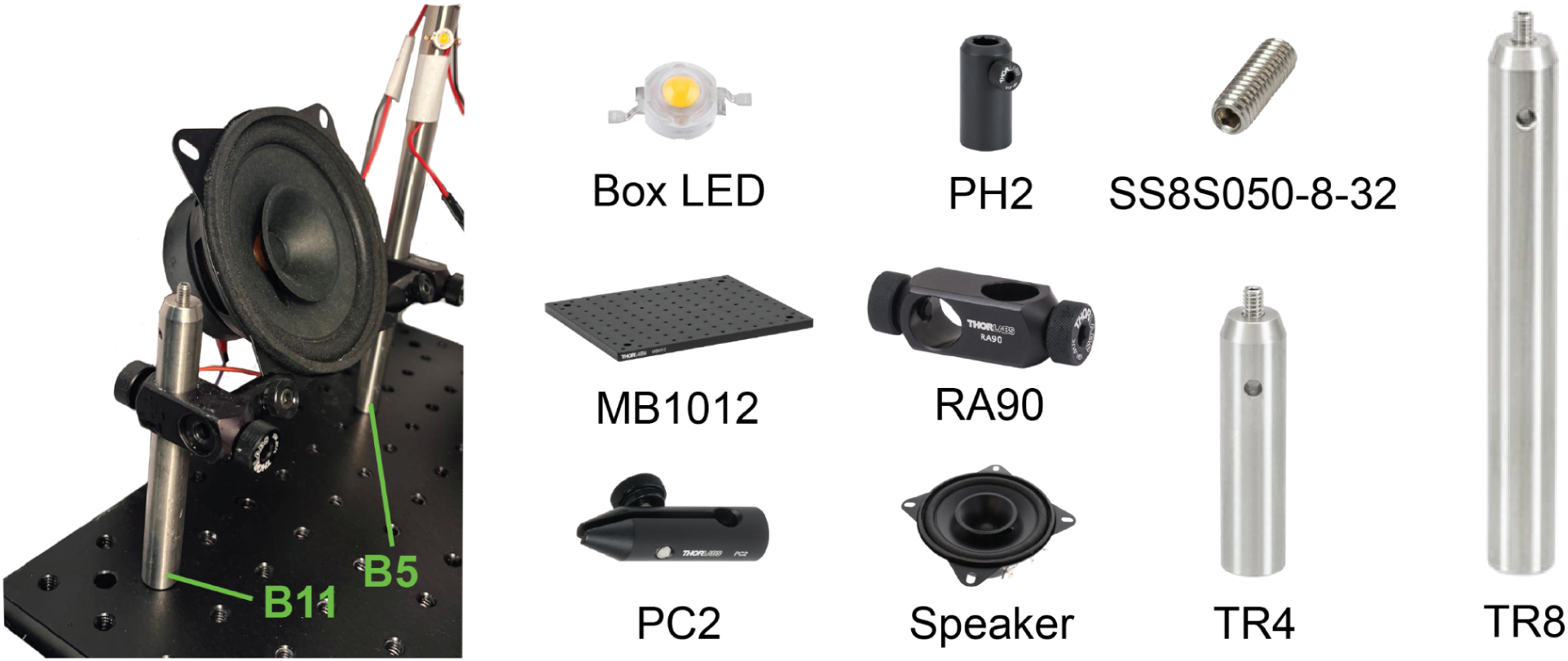
**Speaker assembly**

#### Electronic Diagram

We recommend storing the fully-assembled Arduinos and breadboards at some distance from the behavior box to avoid consequences of spillage of reward solution, escaping mice, etc. In order to allow for easy replacement of components and far reach, we recommend the use of:

a. **Breadboard jumper wires** (e.g., https://www.adafruit.com/product/1953?gad_source=1&gclid=Cj0KCQjwzva1BhD3ARIsA DQuPnX-QaDKtA02VlerAe-222DesWxESo2TQ_uBFPyQyiNimKO1F26yVKgaAhpAEAL w_wcB): soldering these onto the ends of wires of all electrical components used here, e.g., lickometer, speaker, box light, joystick, servo, allows for easy replacement of these components if needed. We also recommend using these as the interface between the breadboard and the stranded wire outlined below as they provide a snug interface between the wire and the breadboard.
b. **Stranded wire** (e.g., https://www.mcmaster.com/8054T13/): allowing some slack of 2-3 meters per electrical component allows for flexibility with Arduino and breadboard placement.

#### Circuit assembly

1. **Prepare the breadboard:** to supply voltage to command and ground strips of the breadboard:

a. connect 5V pin of Arduino 2 to a pin on the command voltage strip of the breadboard, and a ground pin of Arduino 2 to a pin on the ground strip of the breadboard.
b. Pass this potential difference to the strips on the opposite side by connecting a command pin on one side with the command pin on the opposite side, and a ground pin on one side with the ground pin on the opposite side. These connections are shown near the bottom of the breadboard in the figure.
2. **Servo motor:** The servo motor Tower Pro SG5010 (Adafruit: 155) comes with three wires: the brown wire is the ground, the orange wire is the 5V input, and the yellow wire is the input for the pulse-width-modulated (PWM) signal from the Arduino that determines the motor shaft angle.

a. Connect the servo brown wire to a ground pin on Arduino 1.
b. Connect the 5V AC-DC power supply (WSU050-1500, DigiKey: 237-1417-ND) to a barrel-to-jack port adapter (PRT-10288, DigiKey: 1568-1510-ND).
c. Connect the ground terminal of the barrel-to-jack port adapter to a ground pin on Arduino 1
d. Connect the servo orange wire to the positive terminal of the barrel-to-jack port adapter
e. Connect the servo yellow wire to pin 9 on Arduino 2. Note that this pin has a curly line next to it (∼) indicating that it can supply pulse-width-modulated (PWM) input to the servo motor.
3. **Joystick**: The joystick TS6T2S02A (DigiKey: 679-3658-ND) comes with five wires: red and black wires are command voltage and ground wires respectively. Yellow, orange, and blue wires track the analog position of the joystick in three dimensions: anterior-posterior, left-right, and up-down respectively. This setup only tracks anteroposterior (yellow), and up-down displacement (blue) of the joystick. The left-right readout, which is carried by the orange wire, is therefore not used.

a. Connect joystick red wire to the 5V output pin of Arduino 1.
b. Connect joystick black wire to ground pin of Arduino 1.
c. Connect joystick blue wire to pin A1 on Arduino 2.
d. Connect joystick yellow wire to pin A3 on Arduino 2.
4. **Speaker**: Arduino 1 supplies white noise input to the speaker. An arduino cannot generate real white noise, but can rapidly iterate through random tones within a certain range to create the perceptual experience of white noise. This rapid iteration is slowed by any parallel processes (e.g., reading out joystick position, licks, etc.). Therefore, a separate arduino is used for this purpose. To reduce the volume of this output, we use a 160 Ohm resistor in series with the speaker:

a. Connect pin 2 on Arduino 1 to pin 25A on the breadboard.
b. Connect 160 Ohm resistor (DigiKey: 13-MFR-25FRF52-160RTR-ND) to pins 25B and 22B on the breadboard.
c. Connect pin 22A on the breadboard to the positive end of the speaker.
d. Connect the negative end of the speaker to a ground pin on the left side of the breadboard.
e. To control Arduino 1 via Arduino 2, connect Arduino 2 pin 31 to Arduino 1 pin 10, and Arduino 2 pin 33 to Arduino 1 pin 11.
5. **Box light**: Arduino 2 toggles the box light on and off via the breadboard.

a. Connect Arduino 2 pin 2 to pin 16B on the breadboard.
b. Connect pin 16A on the breadboard to the positive end of the box light.
c. Connect the negative end of the box light to the ground strip on the breadboard.
6. **Solenoid valve (“pump”)**: this is powered via a motor driver (SN754410NE, DigiKey 296-9911-5-ND). This motor driver has the capacity to power two solenoid valves or other motors, but we only use its capability for one solenoid in this setup. More details about the functions of each pin in this motor driver can be found in the following instruction manual: https://www.ti.com/lit/ds/symlink/sn754410.pdf?ts=1722601127757&ref_url=https%253A%252F%252Fwww.google.com%252F

a. Insert the motor driver so that the left pins go into pin holes E1-8, and the right pins go into pin holes F1-8. Ensure the divet at the top of the motor driver is oriented toward the top of the breadboard.
b. Connect heat sink (GROUND) pins on the left and right of the motor driver (D4-5 and G4-5) to ground strips on the right and left of the breadboard.
c. Connect pin hole G2 to the command voltage strip of the breadboard on the right, in order to supply 5V power to pin VCC1 of the motor driver to power internal logic translation.
d. Connect the 12V AC-DC power supply (SmoTecQ SA-0243-1202000US, DigiKey: 19262-ND) to a barrel-to-jack port adapter (PRT-10288, DigiKey: 1568-1510-ND).
e. Connect the positive terminal of the barrel-to-jack port adapter to pin D8 on the breadboard (VCC2) to supply the command voltage for the solenoid(s).
f. Connect a 100nF capacitor from pin A8 to the ground strip on the left side of the breadboard.
g. Connect the ground terminal of the barrel-to-jack port adapter to the ground strip on the left side of the breadboard.
h. Connect pin 13 on Arduino 2 to pin D1 on the breadboard (VEN1,2) to enable the channels that will power the solenoid.
i. Connect pin 7 on Arduino 2 to pin D2 on the breadboard (1A). Connect pin 8 on Arduino 2 to pin D7 on the breadboard (2A). These are the driver inputs.
j. Connect pin D3 on the breadboard (1Y) to one terminal of the solenoid, and pin D6 on the breadboard (2Y) to another terminal of the solenoid. The orientation does not matter for the solenoid we used (LHDA1231115H).
7. **Lickometer:** we use an optical lickometer (Sanworks: 1020) that detects breaks in an infrared beam that is shone perpendicularly in front of the lickspout. The setup contains an infrared photoemitter, infrared photodetector, and a built-in LED in the light of the mouse that we use as a Go cue.

a. **Photodetector:**

i. Connect the right command voltage strip to pin J9 on the breadboard.
ii. Connect a 10 kiloOhm resistor (DigiKey: 13-MFR25SFRF52-10KTR-ND) to pins F9 and F12 on the breadboard.
iii. Connect the positive terminal of the infrared photodetector to pin I12 on the breadboard, and the negative terminal of the infrared photodetector to the ground strip on the right side of the breadboard.
iv. Connect Arduino 2 pin 12 to pin J12 on the breadboard. We will use this wire to track whether or not the infrared beam is interrupted by a lick.
b. **Photoemitter**
c. Connect the right command voltage strip to pin J10 on the breadboard.
d. Connect a 330 Ohm resistor (DigiKey: 13-MFR-25FTF52-330RCT-ND) to pins G10 and G13 on the breadboard.
e. Connect the positive terminal of the infrared photoemitter to pin G13 on the breadboard, and the negative terminal of the infrared photoemitter to the ground strip.
f. **Lickometer LED/Go cue light**

i. Connect Arduino 2 pin 10 to pin I11 on the breadboard.
ii. Connect a 1 kiloOhm resistor (DigiKey: 13-MFR-25FRF52-1KTR-ND) to pins H11 and H14 on the breadboard.
iii. Connect the positive terminal of the lickometer LED to pin I14 on the breadboard, and the negative terminal of the lickometer LED to the ground strip on the right side of the breadboard.
8. **Lick indicator LED:** To track in real time whether a mouse is licking from outside of the box, it can be helpful to install a lick indicator LED that shines whenever the infrared beam of the lickometer is broken. To do this:

a. Connect Arduino 2 pin 11 to breadboard pin I15.
b. Connect the positive terminal of the lick indicator LED to pin J15 on the breadboard, and the negative terminal of the lick indicator LED to the ground strip of the breadboard. Any LED will work if it can take an input voltage of 5V; otherwise resistors/different voltage input might be necessary.
9. **Compatibility with fiber photometry and optogenetics.** Our in-house codes use pins 22 and 23 as output TTLs to our photometry system to allow for synchronization of behavior clock with the photometry system clock, and pin 51 to send an output TTL to toggle the on-off state of our optogenetics light sources.

**Figure 9.**
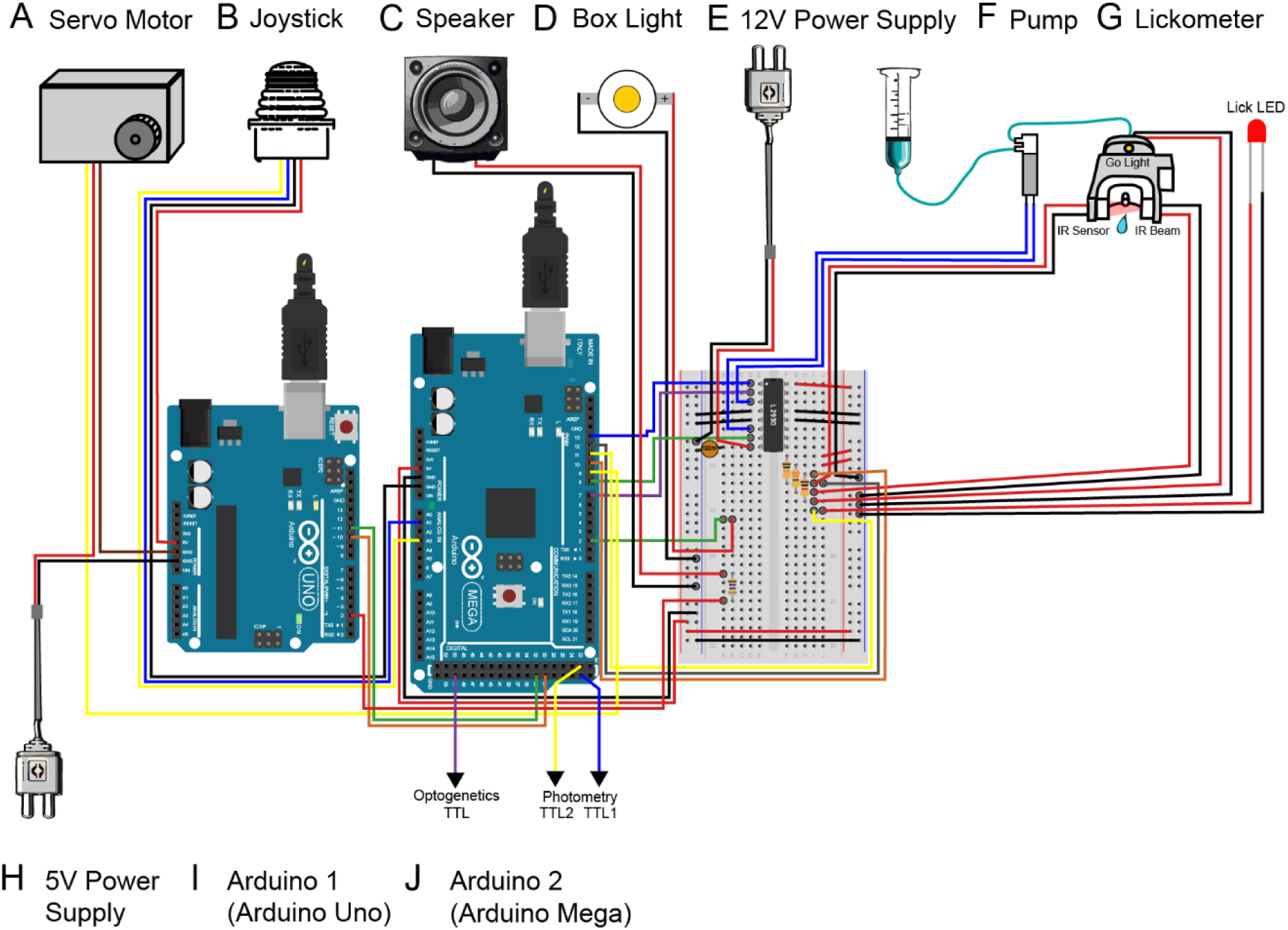
**Value-Based Decision-Making Task electronic diagram**

---

**Table 1.**
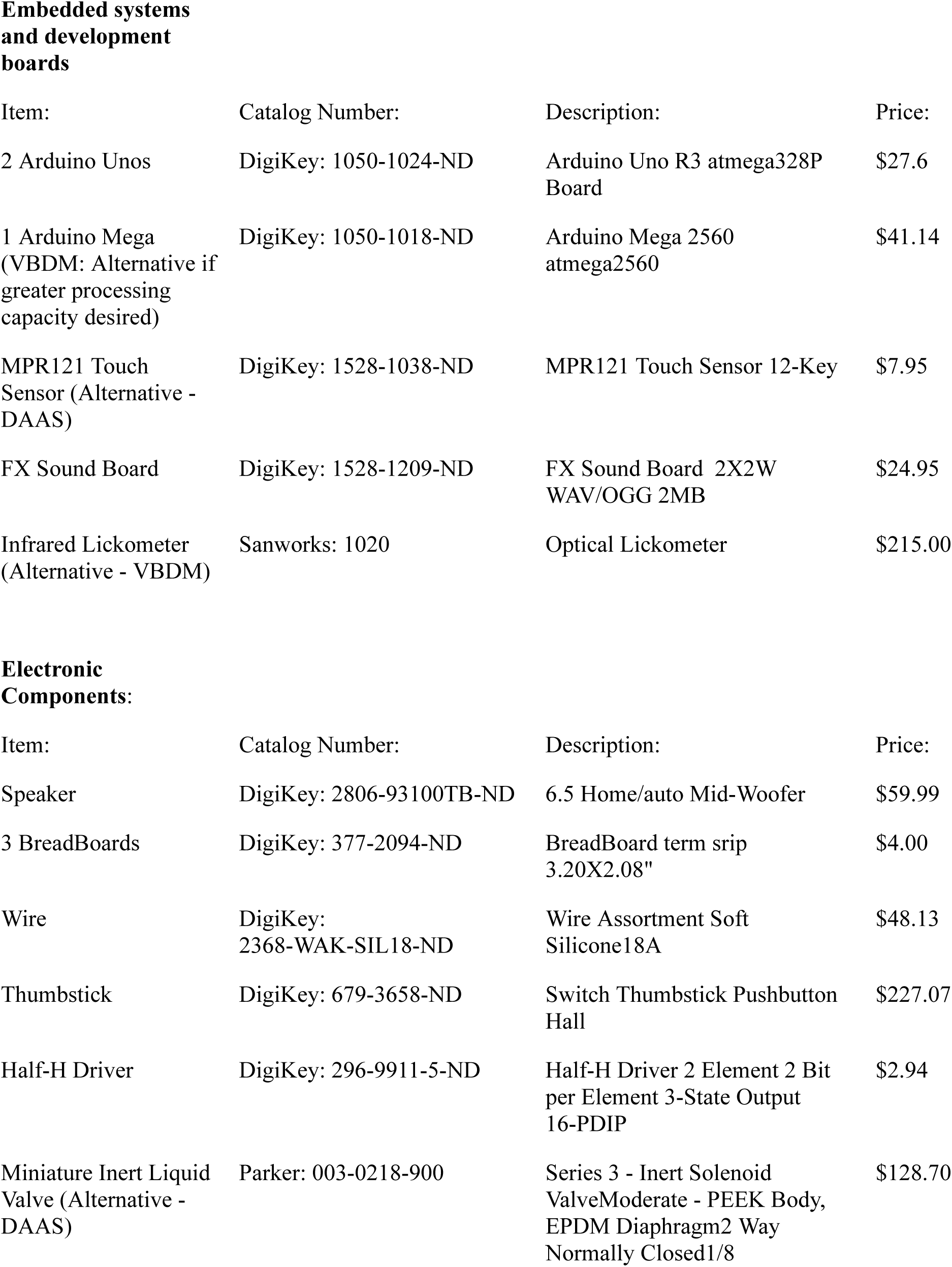
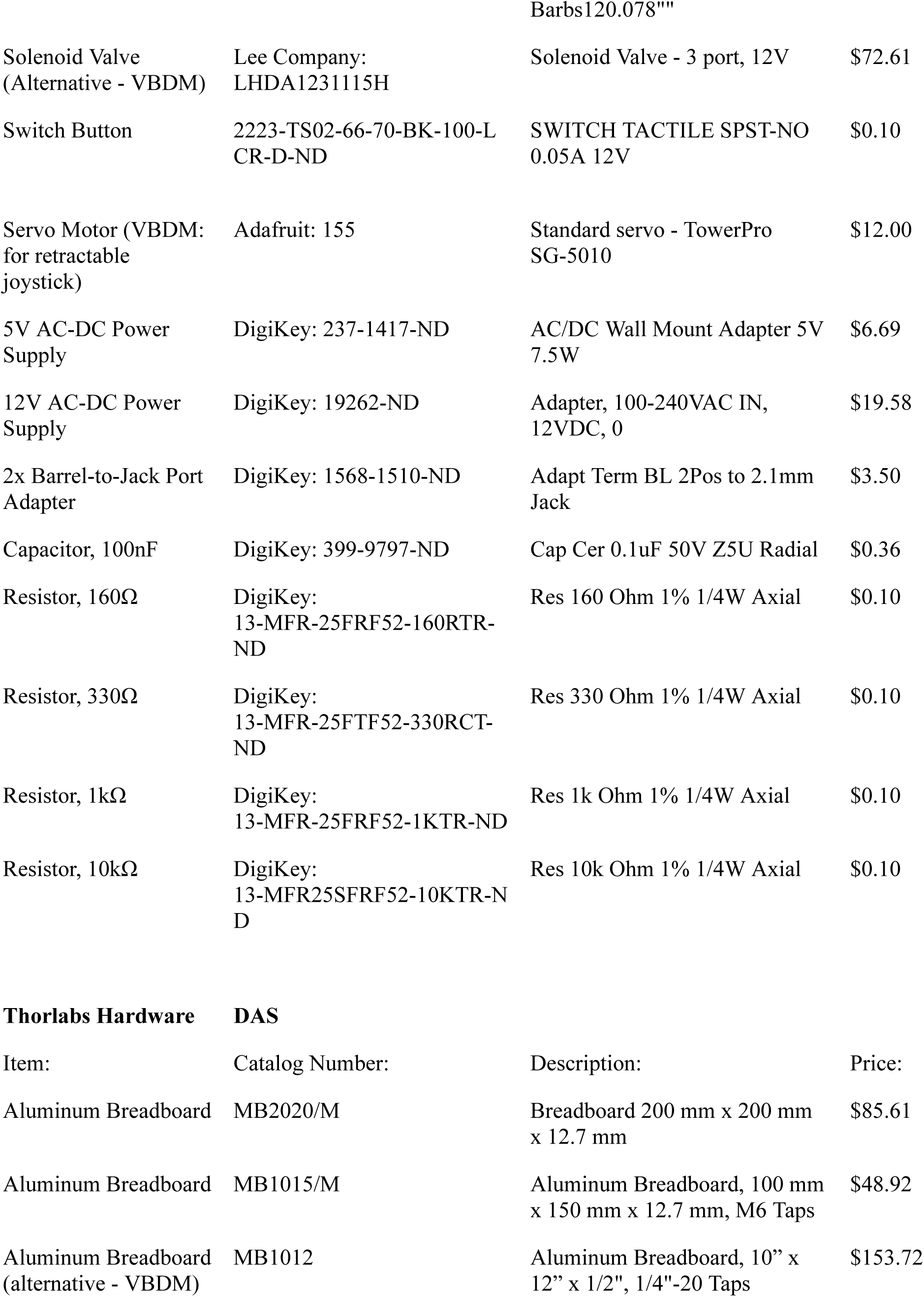
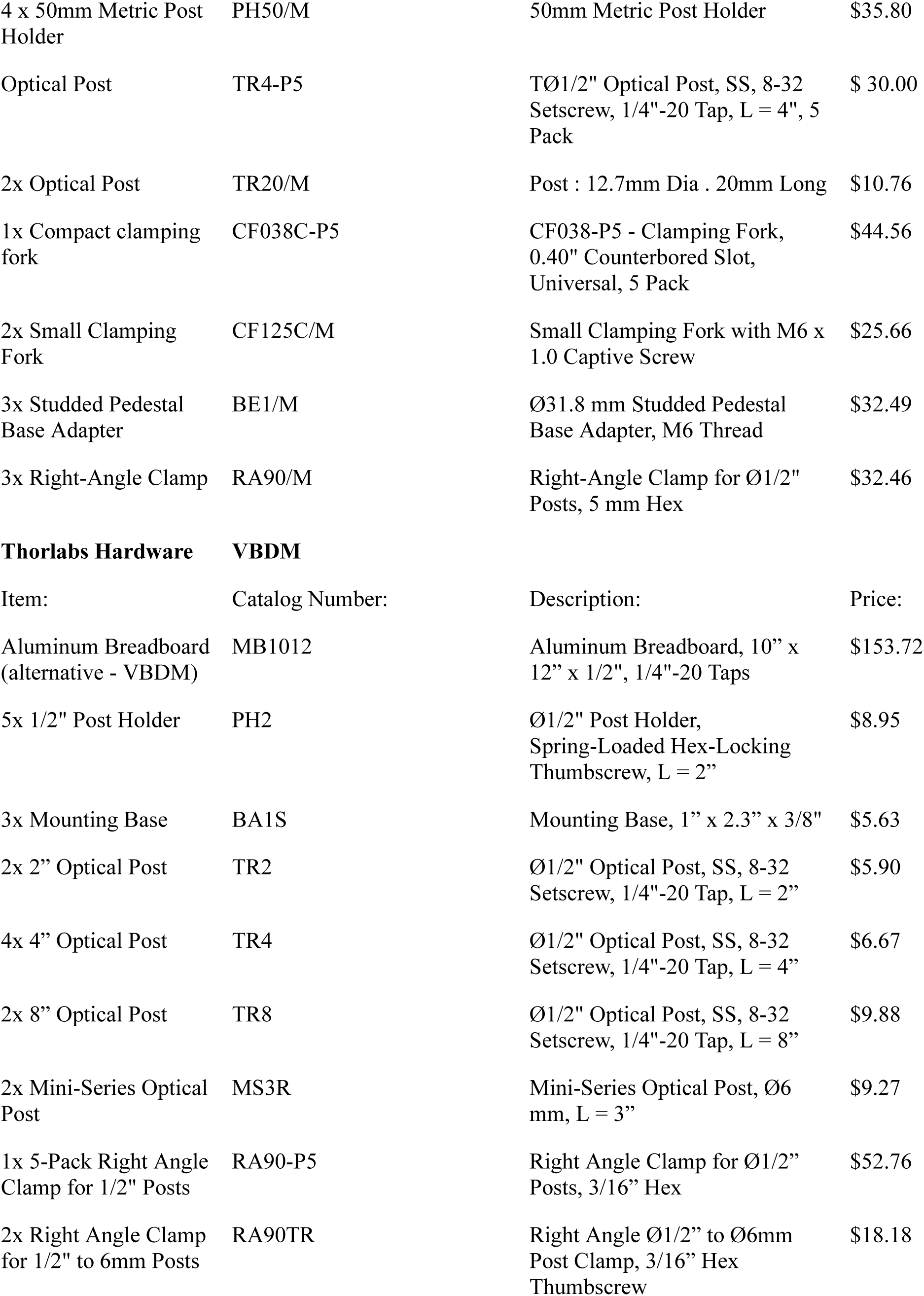
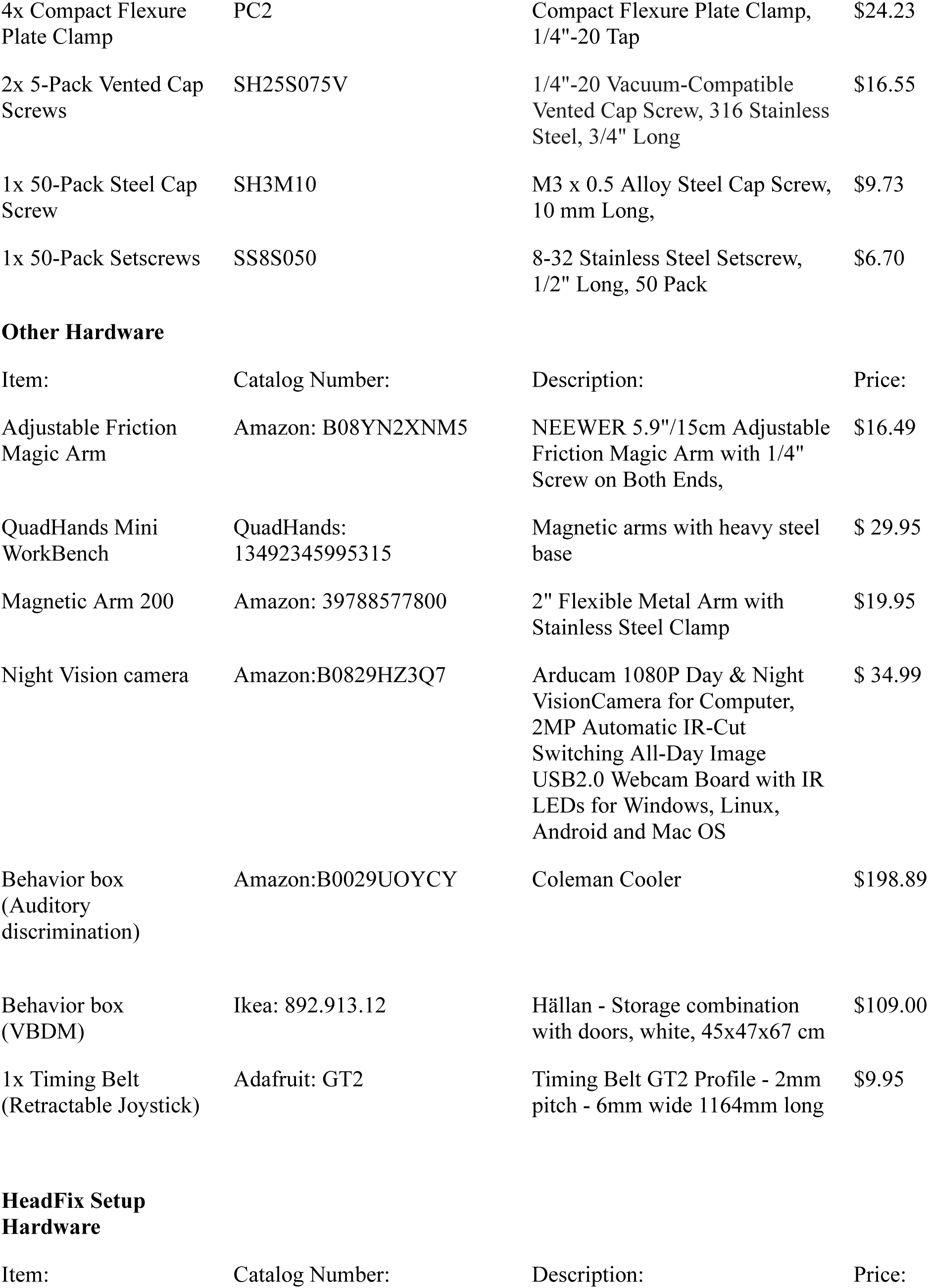
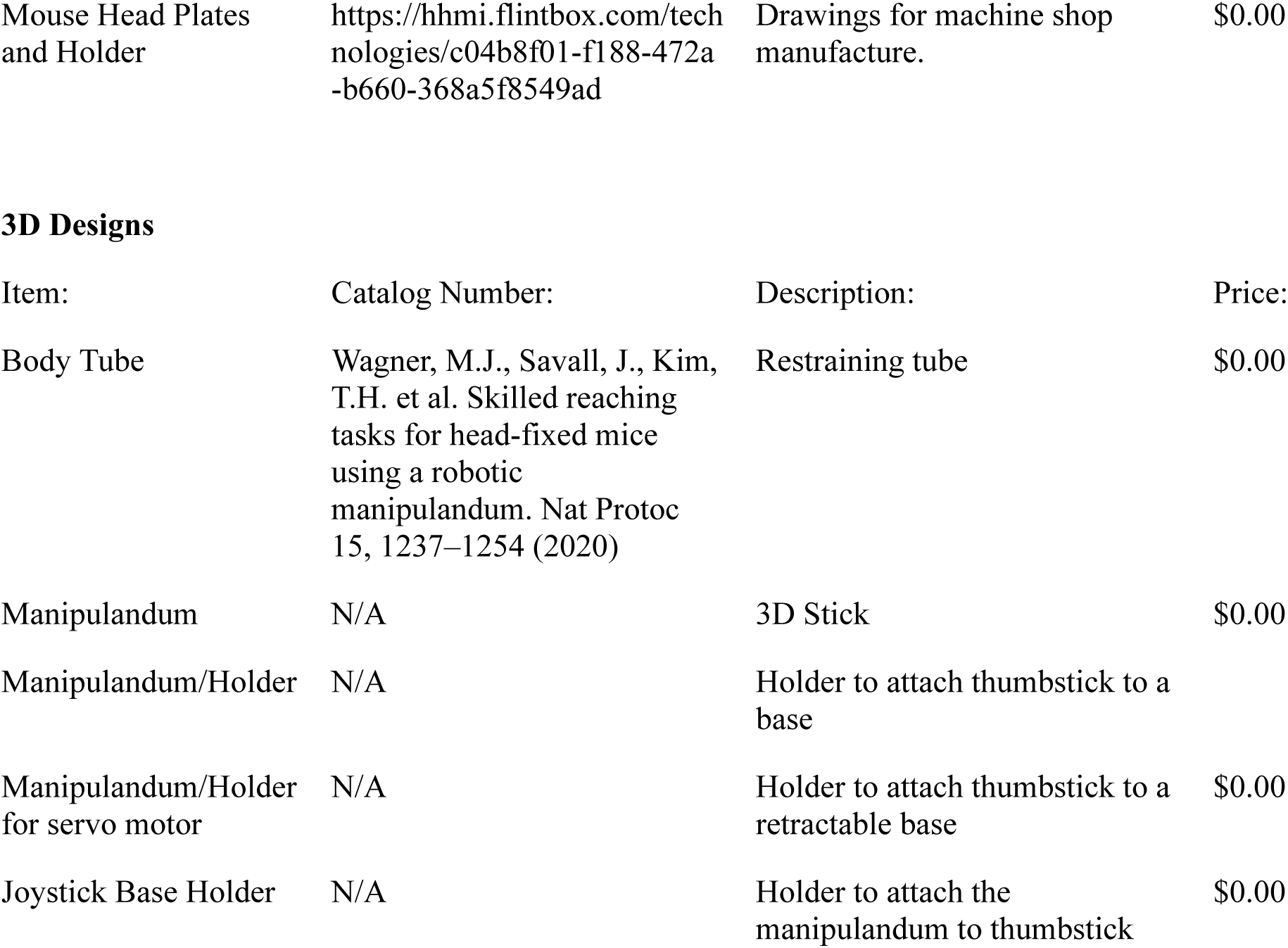

